# Computational design of bifaceted protein nanomaterials

**DOI:** 10.1101/2024.10.18.619149

**Authors:** Sanela Rankovic, Kenneth D. Carr, Justin Decarreau, Rebecca Skotheim, Ryan D. Kibler, Sebastian Ols, Sangmin Lee, Jung-Ho Chun, Marti R. Tooley, Justas Dauparas, Helen E. Eisenach, Matthias Glögl, Connor Weidle, Andrew J. Borst, David Baker, Neil P. King

**Author notes:** Correspondence should be addressed to N.P.K.

## Abstract

Recent advances in computational methods have led to considerable progress in the design of self-assembling protein nanoparticles. However, nearly all nanoparticles designed to date exhibit strict point group symmetry, with each subunit occupying an identical, symmetrically related environment. This limits the structural diversity that can be achieved and precludes anisotropic functionalization. Here, we describe a general computational strategy for designing multi-component bifaceted protein nanomaterials with two distinctly addressable sides. The method centers on docking pseudosymmetric heterooligomeric building blocks in architectures with dihedral symmetry and designing an asymmetric protein-protein interface between them. We used this approach to obtain an initial 30-subunit assembly with pseudo-D5 symmetry, and then generated an additional 15 variants in which we controllably altered the size and morphology of the bifaceted nanoparticles by designing *de novo* extensions to one of the subunits. Functionalization of the two distinct faces of the nanoparticles with *de novo* protein minibinders enabled specific colocalization of two populations of polystyrene microparticles coated with target protein receptors. The ability to accurately design anisotropic protein nanomaterials with precisely tunable structures and functions could be broadly useful in applications that require colocalizing two or more distinct target moieties.

## Introduction

Multi-subunit protein complexes are fundamental for nearly all biological processes and have inspired efforts to design new self-assembling proteins^1–4^. Computationally designed protein nanoparticles have emerged as a promising class of nanomaterials that have served as robust scaffolds for a number of applications, encompassing multivalent antigen presentation^5–14^, structure determination^15–17^, enzyme colocalization^18^, and enhancement of receptor-mediated signaling and virus neutralization^19–21^. Most of the computationally designed nanoparticles described to date have been constructed using a dock-and-design approach in which symmetric building blocks are docked together in a target architecture, and then low-energy protein-protein interfaces are designed between the building blocks to drive assembly^5,22–25^. This approach is efficient in that it allows the construction of large structures from a small number of subunits and minimizes the number of novel protein-protein interfaces that must be designed^3,26^. However, it also constrains the sizes, shapes, and symmetries of the assemblies that can be designed, limiting their functionalization and application. Recent methodological advances have led to the emergence of a novel approach based on the design of pseudosymmetric heterooligomers in which subunits with identical backbone structures but distinct amino acid sequences are arranged symmetrically (e.g., as trimers)^27^. These building blocks allowed extension of the dock-and-design approach to the construction of very large pseudosymmetric protein nanoparticles through the incorporation of several designed protein-protein interfaces in a single material^28,29^. Nonetheless, even these pseudosymmetric nanoparticles have global tetrahedral, octahedral, or icosahedral point group symmetry and, as a result, isotropically distributed subunits across their entire surfaces. Moving beyond isotropic materials to those with controllable anisotropy or directionality requires design methods that incorporate additional asymmetry.

Bifaceted or Janus-like architectures are one such class of anisotropic nanomaterials^30^. Their defining feature is two distinct faces composed of unique molecules that are independently addressable. This property makes Janus-like particles particularly useful in applications that require bringing two different entities together. For example, bispecific T-cell engagers (BiTEs) are a simple and clinically relevant class of such molecules in which a genetic fusion of two different single-chain variable fragments, one against a T-cell marker and the other against a tumor-associated antigen, enhances the anti-cancer activity of T cells by colocalizing them with tumor cells^31^. Self-assembling bifaceted materials have been constructed from a wide range of materials, including metals, silicon dioxide, titanium dioxide, graphene, polyethylene, polystyrene, polyacrylic acid, lipids, and DNA^32–34^. Notably, DNA nanotechnology has been used to construct Janus-like particles that improve cancer vaccination and enhance endosomal escape^33,35^. The design of bifaceted nanoparticles from protein building blocks would be particularly useful due to their multivalency, biocompatibility, modular functionalization through genetic fusion or conjugation of functional domains, and the potential to controllably alter their structures with atom-level accuracy. Despite this potential, methods for accurately designing bifaceted protein nanomaterials have not yet been developed.

Here we develop a general computational approach that enables the design of pseudosymmetric bifaceted protein nanoparticles with precisely tunable structures. We found that displaying protein minibinders on the opposing faces of one such structure enabled it to specifically colocalize polystyrene microparticles coated with therapeutically relevant target proteins, highlighting the potential biomedical utility of this novel class of self-assembling proteins.

## Results

### Computational design of bifaceted pseudo-D5 protein nanomaterials

Designing isotropic protein nanomaterials from oligomeric building blocks minimally requires only a single designed protein-protein interface (**Fig. 1a**), a principle that has been previously leveraged to design a wide variety of novel self-assembling proteins^22,26,36,37^. By contrast, multiple distinct protein-protein interfaces are required to construct anisotropic and bifaceted protein complexes. For example, anisotropic assemblies with D5 symmetry could be constructed from heterotrimeric building blocks by combining i) an asymmetric interface that gives rise to global five-fold rotational symmetry with ii) a symmetric interface along the five dihedral symmetry axes (**Fig. 1b**). In this architecture, each of the three subunits of the heterotrimeric building block is genetically distinct and therefore independently addressable, but the two sides of the overall assembly cannot be uniquely addressed because the trimers would self-assemble. However, if the dihedral interface is instead asymmetric, the resultant pseudo-D5 complexes could not assemble until two different heterotrimers are mixed, yielding bifaceted nanomaterials in which each of the six independent subunits could be uniquely functionalized (**Fig. 1c**). We refer to this pseudo-D5 (pD5) architecture as (ABC)_5_-(ABD)_5_ to indicate its bifaceted nature and hierarchical assembly from heterotrimeric building blocks.

**Figure 1.**
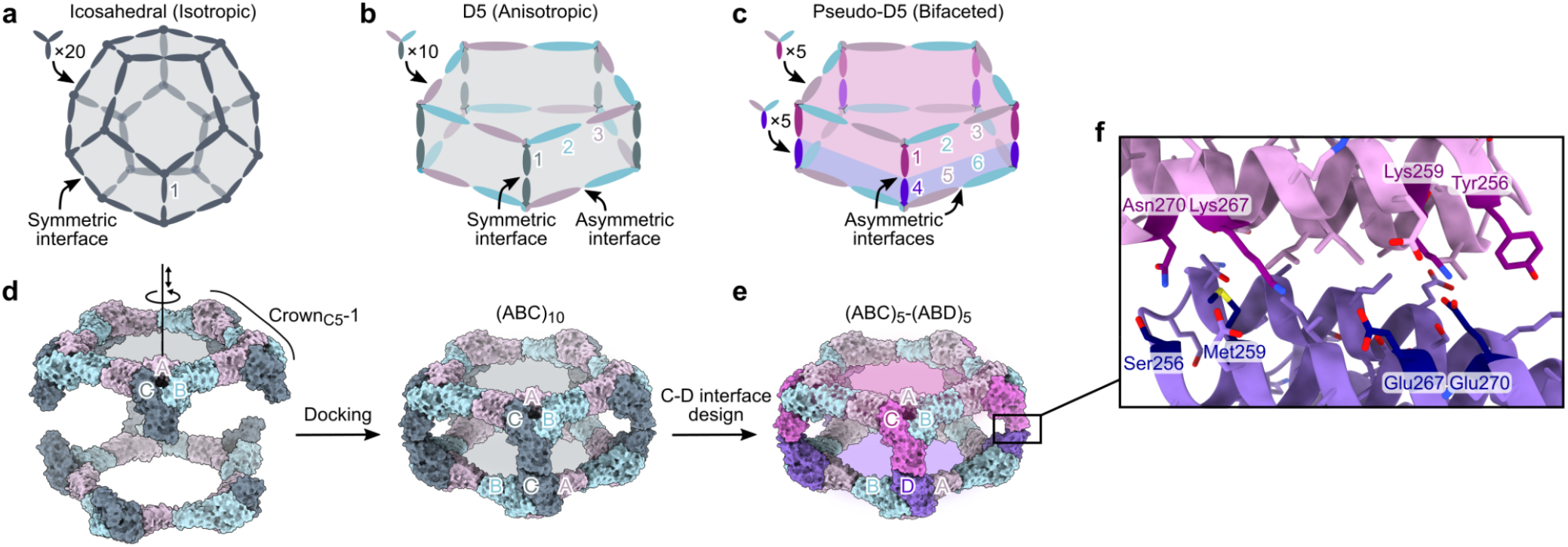
Overview of bifaceted pseudo-D5 architecture and design approach. **a**, A nanoparticle with icosahedral symmetry constructed from homotrimeric building blocks requires only a single (symmetric) designed interface and is isotropic overall. **b**, A nanoparticle with dihedral symmetry (D_*n*_; in this example *n*=5) constructed from a single heterotrimeric building block requires two designed interfaces: an asymmetric interface around the *n*-fold symmetry axis and a symmetric interface along the dihedral two-fold axes. The assembly is anisotropic, but the two opposing faces are constructed from the same three subunits (labeled 1, 2, and 3) and are not independently addressable. **c**, An anisotropic D_*n*_ assembly can be converted to a bifaceted pseudo-D_*n*_ assembly by asymmetrizing the designed interface at the dihedral two-fold axes. The nanoparticle is constructed from two different heterotrimers that can be produced independently prior to nanoparticle assembly, rendering each of the six subunits uniquely addressable, even if subunits 2+6 and 3+5 are genetically identical. **d**, Schematic of the procedure for docking Crown_C5_-1 substructures into (ABC)_10_ assemblies with D5 symmetry. The A, B, and C subunits of Crown_C5_-1 are labeled, and the rotational and translational degrees of freedom sampled during docking are indicated along the five-fold symmetry axis. **e**, An example bifaceted (ABC)_5_-(ABD)_5_ assembly after asymmetric C-D interface design. **f**, Details of an example asymmetric C-D interface. Positions featuring different amino acids in the C and D subunits are highlighted.

We recently showed that stepwise approaches facilitate the design of self-assembling protein complexes featuring multiple designed interfaces^27–29^. To design bifaceted pseudo-D5 nanoparticles, we leveraged a 15-subunit substructure of a recently described pseudosymmetric icosahedral assembly^28^. This ring-like substructure, called Crown_C5_-1, comprises five copies of a pseudosymmetric trimer with three-fold symmetry at the backbone level but distinct amino acid sequences for each subunit (“ABC” heterotrimers). A previously designed interface between the A and B chains drives assembly of the heterotrimers into the 15-subunit substructure with five-fold rotational symmetry, leaving the C-terminal end of the C subunits free for the design of an additional interface. We docked two full-length Crown_C5_-1 assemblies against each other by sampling rotation and translation along the five-fold axis to generate 30-subunit assemblies with D5 symmetry (**Fig. 1d**). To generate additional diversity, we also docked Crown_C5_-1 models in which the C terminus of the C subunit was truncated by up to four α-helices. We then used ProteinMPNN^38^ to design an asymmetric protein-protein interface between the opposing C subunits of each dock to generate sequences intended to form bifaceted (ABC)_5_-(ABD)_5_ complexes (**Fig. 1e**). We employed several negative design strategies during asymmetric interface design by i) biasing ProteinMPNN to favor residues of opposite charge, size, or both on the two sides of the C-D interface, or ii) performing explicit multi-state design to select residues that stabilize the on-target interface and destabilize potential off-target C-C or D-D interfaces (see Methods)^38,39^. These approaches yielded interfaces in which a subset of symmetry-related positions featured distinct amino acids on the C and D subunits (**Fig. 1f**).

We used AlphaFold2 (AF2) structure prediction^40^ of the five C-terminal helices of the C and D subunits to identify sequences strongly predicted to form the on-target C-D interface while not forming off-target C-C and D-D interfaces. Designs for which i) the AF2 prediction of the C-D interface was within 2.0 Å Cα root mean square deviation (RMSD) from the design model, ii) mean predicted aligned error for the inter-chain interactions (mean pAE interaction) was lower than 10, and iii) predicted local distance difference test (pLDDT) was higher than 90 were considered further (**Extended Data Fig. 1a**). Of these, we discarded designs for which structure prediction of either of the off-target C-C or D-D interfaces yielded RMSD < 2.0 Å, mean pAE interaction < 10, and pLDDT > 95 (**Extended Data Fig. 1b**). To further filter for asymmetry-favoring sequences, we identified designs where the difference in mean pAE interaction between the C-D and either C-C or D-D interfaces was greater than 10. Biasing ProteinMPNN to favor residues of opposite charge or opposite charge and size across the C-D interface generated the most designs that passed these criteria. We selected 14 designs from these two groups for experimental characterization, as well as all 7 passing designs from a set in which ProteinMPNN was used with no negative design considerations. The amino acid sequences of all novel proteins generated in this study can be found in **Supplementary Table 1**.

### Experimental characterization of bifaceted pseudo-D5 (pD5) protein nanoparticles

We expressed the A, B, C, and D subunits for each design separately in *E. coli* (with a 6×His tag on the A component), mixed the cells from the A+B+C and A+B+D expression bacterial cultures prior to lysis, and purified the proteins from clarified lysates as cyclic (ABC)_5_ and (ABD)_5_ assemblies using immobilized metal affinity chromatography (IMAC) and size exclusion chromatography (SEC). Despite our negative design strategies, SEC and nsEM showed that nearly all of the designs formed off-target assemblies: 4 yielded either an (ABC)_10_ or (ABD)_10_ assembly, while 16 yielded both (ABC)_10_ and (ABD)_10_ assemblies. These results likely reflect the tendency of even modest protein-protein interactions to drive association of high-symmetry building blocks^41,42^. One design from the set biased to favor residues of opposite charge and size across the C-D interface, called pD5-14, yielded cyclic (ABC)_5_ and (ABD)_5_ assemblies that did not form off-target (ABC)_10_ or (ABD)_10_ assemblies but yielded an earlier elution peak during SEC when mixed, suggesting formation of a larger assembly (**Fig. 2a**). SDS-PAGE of the peak fraction indicated the presence of all four components in the shifted peak (**Fig. 2b**), and dynamic light scattering (DLS) and mass photometry^43^ showed the existence of particles measuring approximately 25.1 nm in size and 1151 kDa in mass, respectively, closely matching the expected values for the target bifaceted nanoparticle (**Extended Data Fig. 2a,b**). Ring-like structures resembling Crown_C5_-1 were observed during nsEM of the pD5-14 (ABC)_5_ and (ABD)_5_ components as expected, along with a substantial fraction of unassembled heterotrimeric building blocks (**Fig. 2c,d**). By contrast, electron micrographs of the purified pD5-14 (ABC)_5_-(ABD)_5_ complexes revealed monodisperse fields of particles that resembled the intended 30-subunit structure, with considerably fewer unassembled heterotrimers (**Fig. 2e**). Rigid-body fits of the pD5-14 (ABC)_5_, (ABD)_5_, and (ABC)_5_-(ABD)_5_ design models into the corresponding low-resolution 3D reconstructions strongly suggested that each assembly adopted the intended structure. The (ABC)_5_-(ABD)_5_ complexes were remarkably thermostable: essentially no changes in hydrodynamic diameter, intrinsic tryptophan fluorescence, or static light scattering were observed while heating pD5-14 complexes from 25–95°C, indicating that the assemblies do not unfold or aggregate even when subjected to near-boiling temperatures (**Extended Data Fig. 2c–e**).

**Figure 2.**
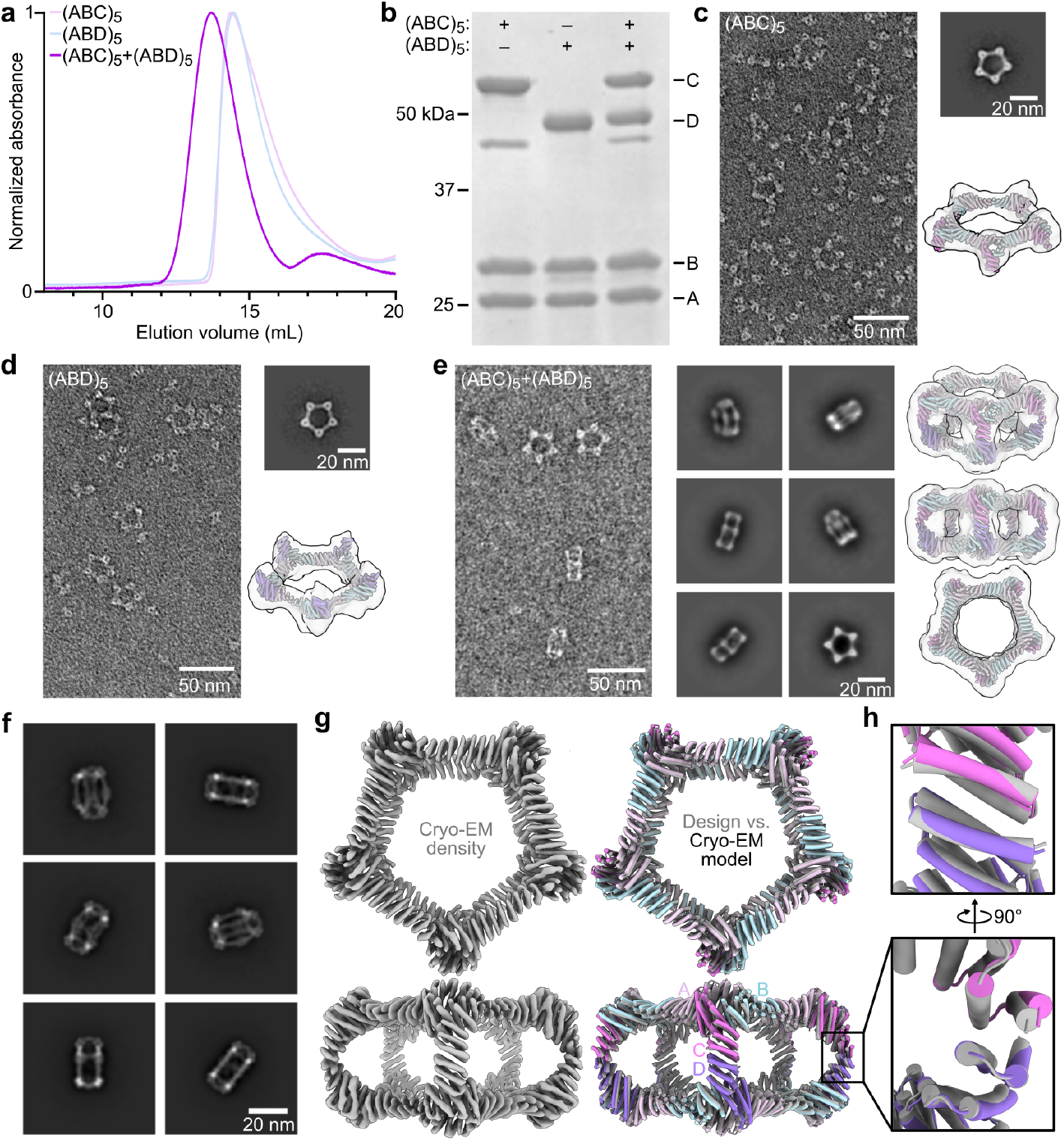
*In vitro* assembly and structural characterization of pD5-14. **a**, SEC traces of pD5-14 (ABC)_5_, (ABD)_5_, and (ABC)_5_-(ABD)_5_ assemblies. **b**, SDS-PAGE of SEC-purified assemblies from panel (**a**). The locations of the A, B, C, and D subunits, as well as those of marker bands, are indicated. **c–e**, nsEM of pD5-14 (ABC)_5_ (**c**), (ABD)_5_ (**d**), and (ABC)_5_-(ABD)_5_ (**e**) assemblies, including representative 2D class averages and rigid-body fits of the computational design models into 3D reconstructions. **f**, Cryo-EM 2D class averages of pD5-14 (ABC)_5_-(ABD)_5_. **g**, *Left:* 4.30 Å resolution cryo-EM density map of pD5-14 (ABC)_5_-(ABD)_5_ viewed along two orthogonal axes. *Right:* Overlay between the computational design model (gray) and the experimentally determined cryo-EM model (colors). **h**, Details of asymmetrically designed C-D interface. Scale bars: 50 nm (raw micrographs) and 20 nm (2D class averages).

To evaluate the accuracy of our design approach at higher resolution, the pD5-14 (ABC)_5_-(ABD)_5_ sample was vitrified and imaged using cryo-electron microscopy (cryo-EM; **Extended Data Fig. 3a**). 2D class averages clearly indicated the formation of particles with the intended morphology (**Fig. 2f** and **Extended Data Fig. 3b**), and a single-particle reconstruction using C5 symmetry resulted in a 4.30 Å volume map in which individual alpha helices were well resolved (**Fig. 2g** and **Extended Data Fig. 3c–f**). Relaxing the helices of the pD5-14 design model into the cryo-EM density resulted in a structure that matched the design model remarkably well, yielding a backbone RMSD of 3.0 Å over the entire 30-subunit assembly. Aligning only the two subunits comprising the asymmetric C-D interface yielded a backbone RMSD of 1.3 Å, although we could not distinguish the C and D chains from each other since side chains were not resolved (**Fig. 2h**). Overall, our cryo-EM analysis of pD5-14 (ABC)_5_-(ABD)_5_ complexes assembled *in vitro* confirmed that our approach enables the design of bifaceted protein nanomaterials with high accuracy.

### Fine-tuning of bifaceted protein nanoparticle size and shape using RFdiffusion

Many biological phenomena, including T-cell activation, synaptic transmission, and exocytosis, strongly depend on the distance between two biological objects such as cells or secretory vesicles^44–46^. However, methods or tools for bringing two entities together at prescribed distances are still lacking. We leveraged the recently developed machine learning-based design tools RFdiffusion^47^ and ProteinMPNN^38^ to design bifaceted nanomaterials with systematically varying size and shape by generating *de novo* extensions within the C subunit of pD5-14. Specifically, we “cut” the loop preceding the last two helices of the C subunit to leave the asymmetric C-D interface intact and translated the remainder of the (ABC)_5_ substructure 25, 50, 75 and 100 Å along the five-fold axis (**Fig. 3a**). We defined an additional target in which we translated the (ABC)_5_ substructure 50 Å and also rotated it 25°. For each target structure, we used RFdiffusion to generate *de novo* protein backbone connecting the translated portion of the C subunit to its interface-forming C-terminal helices, designed amino acid sequences for the *de novo* extensions using ProteinMPNN, and predicted the structures of the designed C subunits using AF2 (**Fig. 3b**). The number of residues making up the *de novo* extensions in designs that passed our AF2 filters (pLDDT > 90 and RMSD to the RFdiffusion output < 1.5 Å) differed between each target architecture (**Fig. 3c**). For example, passing designs for the 25 Å extension were narrowly distributed around 75 inserted amino acids, whereas 300–400 inserted amino acids were required for 100 Å extensions. The *de novo* backbones generated by RFdiffusion were often helical repeats that resembled the topology of the original subunit, and inserting more diffused residues for a given extension generally led to an additional alpha helix or occasionally extruded loops (**Extended Data Fig. 4a**). Interestingly, a higher proportion of backbones comprising beta sheets was observed for the 75 and 100 Å extensions, but none of these designs passed our AF2 filters (**Extended Data Fig. 4b**). We selected for experimental characterization the designs with the lowest RMSDs between their AF2 predictions and diffused backbones, comprising 7 designs for the 25 Å extension and 10 each for the 50, 75, and 100 Å extensions as well as the 50 Å extension with a 25° rotation.

**Figure 3.**
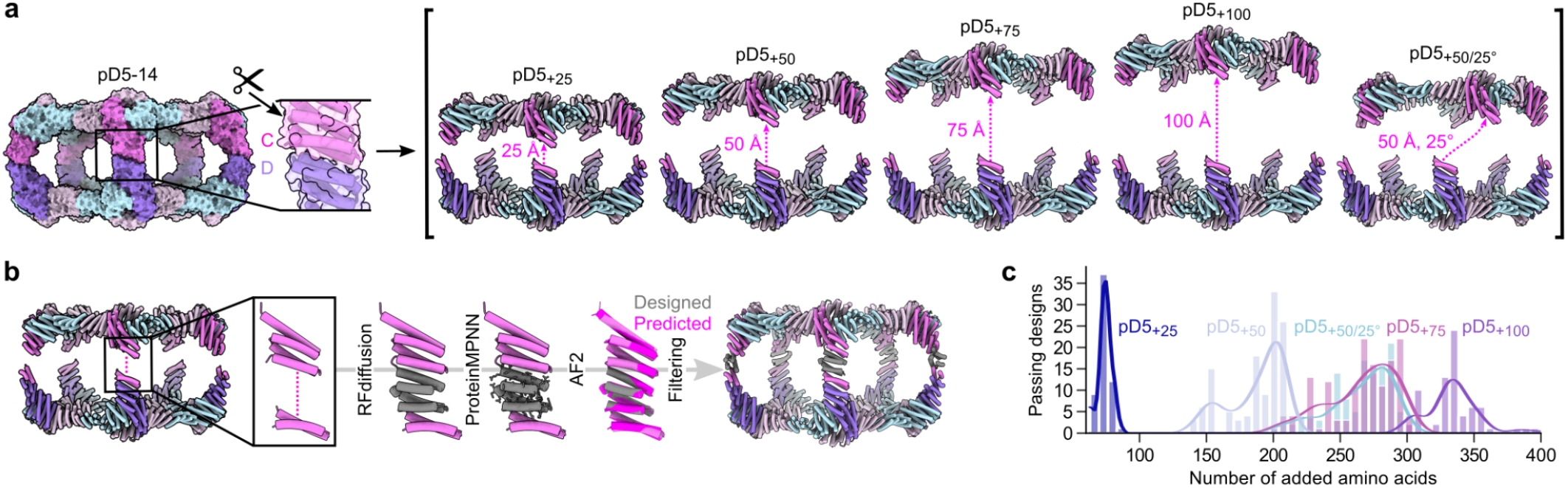
Fine-tuning of bifaceted protein nanoparticle size and shape using RFdiffusion. **a**, Schematic of the approach used to define target extended architectures. *Left:* The scissors indicate the loop in the C subunit that was “cut”. *Right:* Target architectures were defined by translating and in one case rotating most of the (ABC)_5_ substructure while leaving the designed asymmetric C-D interface intact. **b**, Schematic of the design process used to generate *de novo* C subunits for each extended architecture. **c**, Number of designs that passed AF2 filtering (pLDDT > 90; RMSD to design model < 1.5 Å) per number of added amino acid residues for each extended architecture.

We expressed and purified each new (ABC)_5_ design as described above, mixed them with pD5-14 (ABD)_5_, and assessed pD5 particle formation by SEC (**Extended Data Fig. 5, 6**). We observed peaks at the expected elution volumes for two to four of the designs from each of the extended architectures (**Supplementary Table 1**). The SEC traces also indicated that in all cases a substantial portion of each mixture remained as unassembled (ABC)_5_ and (ABD)_5_ components. Three of the four successful designs extended by 100 Å were derived from the same RFdiffusion-generated backbone and differed only at the sequence level (pD5_+100_-46, pD5_+100_-48, and pD5_+100_-71). As with pD5-14, nsEM showed that each extended (ABC)_5_ component prior to mixing with (ABD)_5_ formed only ring-like substructures resembling Crown_C5_-1, with substantial amounts of unassembled trimer also present (**Fig. 4** and **Extended Data Fig. 6**). By contrast, after mixing with pD5-14 (ABD)_5_, D5-like assemblies with aspect ratios clearly different from pD5-14 were the predominant species, with some unassembled trimer present as well. While 2D class averages of particles extended by 25, 50, 75, and 100 Å each strongly resembled the five-pointed star of pD5-14 viewed along its five-fold symmetry axis, the averages of pD5_+50/25°_-344 uniquely showed two lobes at each of the five points, consistent with the designed 25° rotation. Low-resolution single-particle reconstructions of one assembly of each type confirmed that they adopted the intended structures, including the “bent” pillars connecting the two distinct faces of pD5_+50/25°_-344. These data establish that our computational approach can accurately generate bifaceted protein nanomaterials with precisely tunable structures.

**Figure 4.**
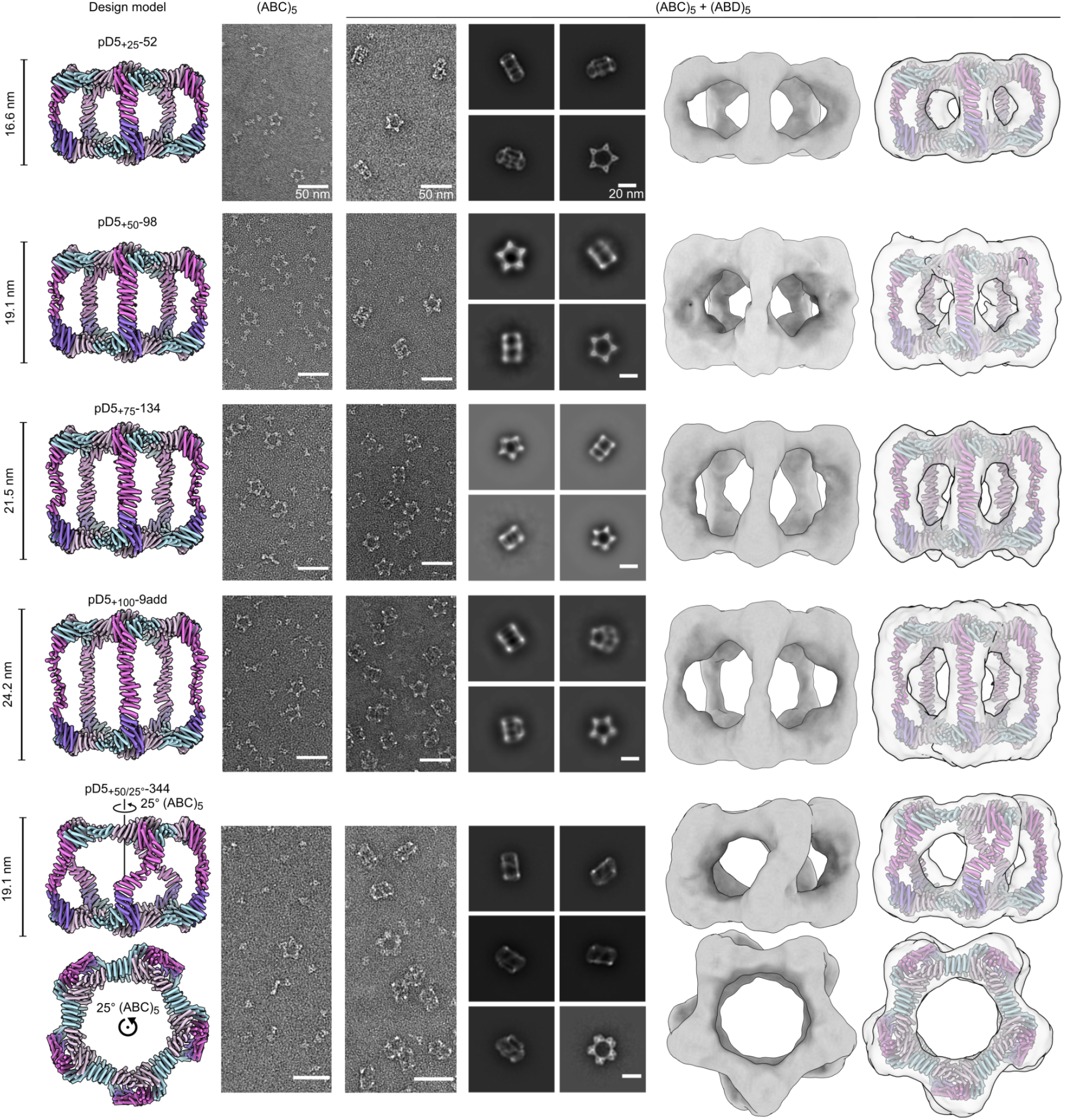
nsEM characterization of extended (ABC)_5_-(ABD)_5_ nanoparticles. *From top to bottom:* Representative assemblies extended by 25, 50, 75, and 100 Å, or extended by 50 Å and rotated by 25°. *From left to right:* computational design models, raw micrographs of (ABC)_5_, raw micrographs of (ABC)_5_+(ABD)_5_, 2D class averages of (ABC)_5_+(ABD)_5_ showing representative top and side views, 3D reconstructions, and design models fit into the 3D reconstructions. Side views of the design models and 3D reconstructions are shown for each particle except pD5_+50/25°_-344, for which side and top views are shown. Scale bars: 50 nm (raw micrographs) and 20 nm (2D class averages).

### Minibinder-functionalized bifaceted nanoparticles colocalize two distinct fluorescent polystyrene microparticle populations

To demonstrate the ability of the bifaceted nanoparticles to colocalize two distinct entities, we sought to genetically fuse different *de novo* protein minibinders^48,49^ to the subunits making up each face of pD5-14. However, neither terminus of any subunit was initially available for genetic fusion, as the C termini were all involved in the protein-protein interfaces that drive bifaceted nanoparticle assembly and the N termini were oriented toward the nanoparticle interior. We therefore redesigned the first three helices in all subunits of pD5-14—which were identical at the backbone level—to generate new, outward-facing N termini. Specifically, we employed block adjacency matrices in RFdiffusion, which enable precise definition of target topologies and contacts^47^, to i) change the order of the first three helices, ii) insert a new alpha helix to effect the desired change in directionality, and iii) construct new loops between the redesigned helices (**Fig. 5a**). We generated 100 novel protein backbones, designed sequences for them using ProteinMPNN, and filtered the designs using AF2 prediction, selecting for experimental characterization the ten designs with pLDTT > 90 and the lowest predicted RMSD to the original heterotrimer.

**Figure 5.**
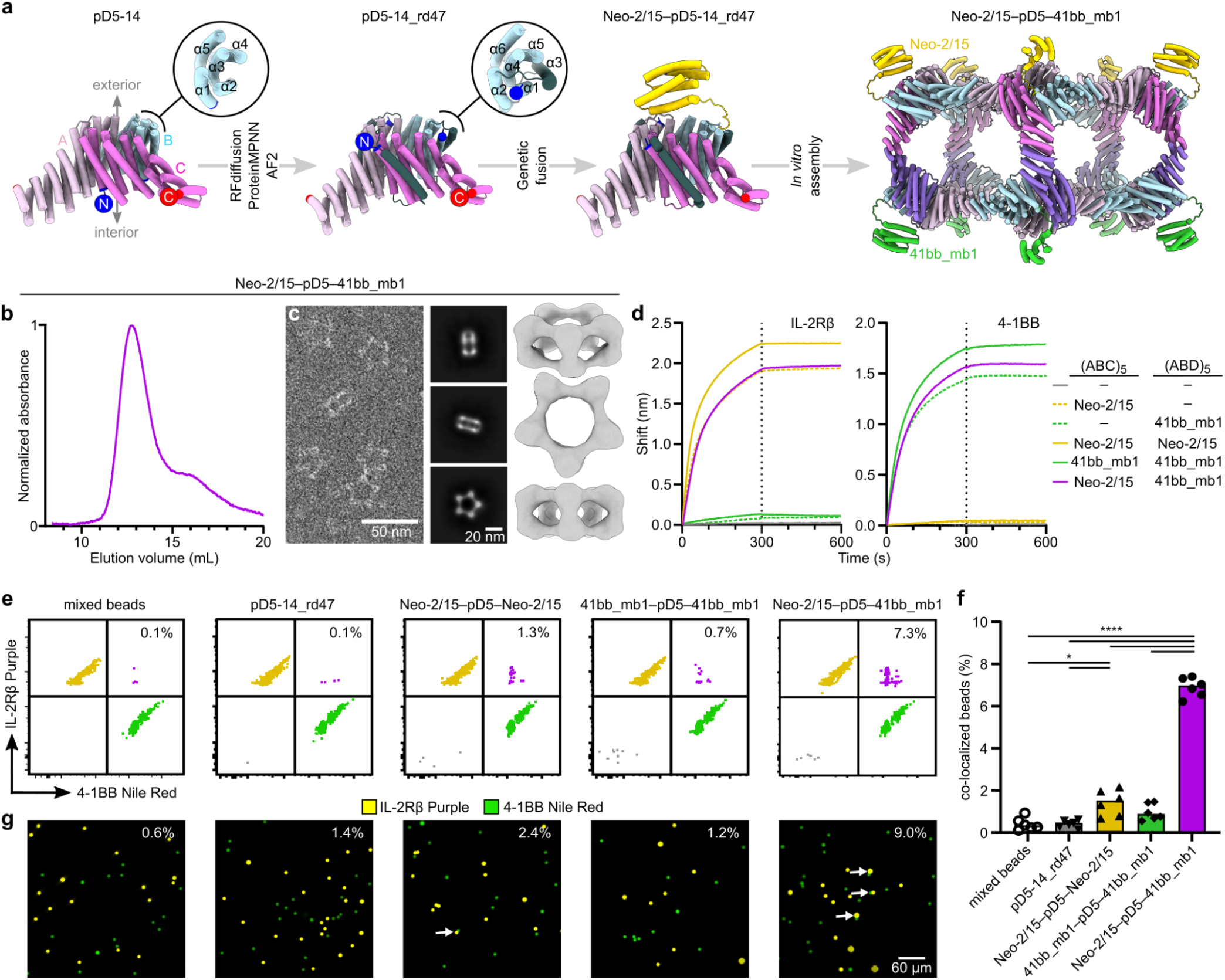
Minibinder-functionalized bifaceted nanoparticles colocalize two distinct fluorescent polystyrene microparticle populations. **a**, Schematic of the approach used to i) reorient the N termini of the A, B, and C subunits of pD5-14 to face toward the nanoparticle exterior and ii) independently functionalize each face of the resultant bifaceted nanoparticle with protein minibinders. The newly diffused α-helix is shown in dark gray, and N and C termini are indicated by blue and red circles, respectively. **b**,**c**, SEC trace (**b**) and nsEM (**c**) of pD5-14_rd47 with Neo-2/15 and 41bb_mb1 genetically fused to the B components of the (ABC)_5_ and (ABD)_5_ components, respectively. Scale bars: 50 nm (raw micrograph) and 20 nm (2D class averages). **d**, Binding of functionalized bifaceted nanoparticles to IL-2Rβ (left) and 4-1BB (right) ectodomains, measured by BLI. The legend indicates which minibinder is fused to each component in each bifaceted nanoparticle. **e**, Representative flow cytometry plots showing colocalization of IL-2Rβ–coated Nile Red and 4-1BB–coated Purple Polystyrene particles by functionalized bifaceted nanoparticles. **f**, Quantitation of colocalization detected by flow cytometry. Each point represents an independent measurement, and the height of each bar indicates the mean for each group. Statistical significance was calculated via one-way ANOVA with Geisser-Greenhouse correction followed by Tukey’s multiple comparisons test, with individual variances computed for each comparison. *, p < 0.1; ****, p < 0.0001. **g**, Representative fluorescence microscopy images showing colocalization of IL-2Rβ–coated Nile Red and 4-1BB–coated Purple Polystyrene particles by functionalized bifaceted nanoparticles. The percentage of colocalized beads observed across 25 independent fields of view is displayed in the upper right corner of each image. Scale bar: 60 μm.

Two of the designs, pD5-14_rd47 and pD5-14_rd106, yielded SEC peaks at the retention volume expected for the target bifaceted nanoparticle after mixing the redesigned (ABC)_5_ and (ABD)_5_ components (**Extended Data Figs. 7a**,**b and 8a**,**b**). Analysis of the SEC-purified assemblies by DLS indicated the existence of uniform particles of the expected size (∼25 nm; **Extended Data Figs. 7c and 8c**). As before, nsEM of the separate (ABC)_5_ and (ABD)_5_ components showed only ring-like substructures and unassembled trimers, while analysis of the purified pD5-14_rd47 and pD5-14_rd106 (ABC)_5_-(ABD)_5_ assemblies yielded 2D class averages and 3D reconstructions that matched the size and morphology of the respective bifaceted nanoparticle design models (**Extended Data Figs. 7d and 8d**).

We made genetic fusions of the IL-2Rβ–targeting minibinder Neo-2/15 (ref. ^48^) and the 4-1BB–targeting minibinder 41bb_mb1 (ref. ^50^) to the redesigned pD5-14_rd47 B subunit and purified Neo-2/15–bearing (ABC)_5_ and 41bb_mb1-bearing (ABD)_5_ components as described above. After mixing, SEC and nsEM confirmed the formation of the expected bifaceted nanoparticle, although the displayed minibinders could not be visualized due to their small size and the flexible genetic linker used (**Fig. 5b,c**). The dual-functionalized nanoparticle bound both IL-2Rβ and 4-1BB in biolayer interferometry studies (BLI), while control bifaceted nanoparticles displaying only one of the two minibinders or the same minibinder on both faces (**Extended Data Fig. 9**) only bound the cognate receptor but not the non-cognate receptor (**Fig. 5d**).

We then used the minibinder-displaying pD5-14_rd47 complexes to colocalize two distinct populations of fluorescent polystyrene microparticles. We first coated the microparticles with target receptors by separately conjugating biotinylated 4-1BB and IL-2Rβ to streptavidin-coated Nile Red- and Fluorescent Purple-labeled microparticles, respectively. After mixing the two microparticle populations and incubating them with unfunctionalized (“bare”) pD5-14_rd47, control bifaceted nanoparticles displaying the same binder on both faces, or the bifaceted nanoparticle displaying the two different binders on opposing faces, we measured colocalization using flow cytometry and fluorescence microscopy. Incubation with bare pD5-14_rd47 did not increase colocalization above background levels by flow cytometry (0.1%), while incubation with bifaceted nanoparticles displaying Neo-2/15 or 41bb_mb1 on both sides led to slightly increased numbers of double-positive events (1.3% and 0.7%, respectively; **Fig. 5e,f**). By contrast, incubation with the bi-functionalized nanoparticle significantly increased the number of double-positive events (7.3%), indicating that displaying different protein minibinders on the two faces of pD5-14_rd47 efficiently colocalized the two populations of microparticles. Similar results were obtained by fluorescence microscopy, where 9.0% of the visualized microparticles were colocalized by the bi-functionalized nanoparticles, compared to 1.2–2.4% colocalization after incubation with the three control nanoparticles (**Fig. 5g**). These data establish that our computationally designed bifaceted nanoparticles can be functionalized to colocalize two distinct biological entities.

## Discussion

Our results establish a general computational approach for accurately designing bifaceted protein nanomaterials with customizable structures. Breaking symmetry is a current focus of innovation in the computational design of novel self-assembling proteins, as it simultaneously provides a route to much larger materials and unlocks the ability to address and functionalize specific locations (e.g., subunits). A pair of recent reports from our groups used pseudosymmetric heterooligomeric building blocks^27^ to build very large protein assemblies^39,51^, yet these were still isotropic and therefore had limited addressability. Here we go beyond those strategies by designing anisotropic assemblies with two distinctly addressable faces. We are aware of only one previous report of an engineered multivalent Janus-like protein nanoparticle. In that study, mutations were introduced into a naturally occurring D5 assembly (*Brucella* lumazine synthase) that disfavored pentamer homodimerization and favored pentamer heterodimerization^52^. Here we generated new bifaceted nanoparticle architectures with target structural features by combining protein-protein docking, asymmetric interface design, and generative design of *de novo* subunits. The pseudosymmetric, 30-subunit bifaceted nanoparticles we described do not, to our knowledge, resemble any known naturally occurring or engineered protein complexes. A limitation of the present study is that we only demonstrated the design of assemblies with pseudo-D5 symmetry. However, as demonstrated by the generalization of our previously described dock-and-design approach to a wide variety of symmetric architectures^5,22,23,25,36,37,53,54^, our approach is not limited to pD5 architectures and in principle could be used to design assemblies with pseudo-D2, -D3, or any other dihedral symmetry.

This study also presents, to our knowledge, one of the first methods capable of precisely tuning the structures of designed self-assembling proteins. Most design methods to date have aimed at the more achievable goal of generating single, well-defined target structures due to the sheer complexity of designing novel protein assemblies^5,22–25,37,38,47,53–61^. Nevertheless, designing protein nanomaterials with tunable structures has been a long-standing goal with many potential applications. For example, we exploited the modularity and extensibility of coiled-coils to show that varying antigen-antigen spacing on protein nanoparticle immunogens significantly influences their immunogenicity^62^. Recently, Huddy et al. reported a “copy-paste” approach to precisely and systematically alter the structures of protein assemblies built from twistless repeat protein building blocks^58^. Both of these methods are remarkable for their simplicity, but are strictly limited to specific types of regularly repeating protein building blocks: coiled-coils and twistless repeat proteins, respectively. Here, we took the opposite approach: we achieved precise and systematic control over nanoparticle morphology by leveraging recent advances in AI-based protein structure prediction and design^38,40,47^ to create bespoke subunits of the desired sizes and shapes. We obtained multiple hits for all five extended architectures by experimentally screening ten or fewer candidates, a success rate that compares favorably to those observed historically for designing new self-assembling proteins. Combined with the accompanying manuscript by Wang et al., our results show that this strategy for fine-tuning the structures of self-assembling protein complexes—inpainting between existing protein-protein interfaces arranged in space—should generalize to any symmetric or asymmetric architecture and enable the accurate design of a wide variety of custom self-assembling protein complexes.

We demonstrated the potential utility of the bifaceted nanoparticles by displaying different protein minibinders on each face and using them to specifically colocalize beads coated with two distinct receptor proteins. This required altering the tertiary structure of the nanoparticle subunits so that they had exterior-facing N termini for minibinder display. A similar RFdiffusion-ProteinMPNN-AF2 pipeline, here incorporating block adjacency, again proved successful, yielding two successful designs out of ten tested and demonstrating another level of control over the structure of designed protein assemblies. We note that even though we only displayed minibinders on the B subunits, all six of the subunits in the bifaceted (ABC)_5_-(ABD)_5_ nanoparticles reported here are uniquely addressable. This property derives not only from the well-defined bifaceted architecture of the (ABC)_5_-(ABD)_5_ assemblies, but also from their hierarchical assembly *in vitro*. That is, the separate assembly of the (ABC)_5_ and (ABD)_5_ components enabled the B subunits on each side to be genetically distinct or independently functionalized. We exploited this property to display different protein minibinders on each face and colocalize receptor-coated beads. Together with the precise structural control afforded by our design approach, this proof of principle demonstrates the potential of computationally designed bifaceted nanoparticles to colocalize distinct biological entities at prescribed distances.

## Acknowledgements

We thank Joe Watson and David Juergens for helpful discussions, Quinton Dowling for providing RPXdock scripts, Luki Goldschmidt and Patrick Vecchiatio for maintaining computing resources, Kandise VanWormer and Hernan Nunez-Ortega for wet lab management, and Joel Quispe and Sasha Dickinson for management of the University of Washington cryo-EM facilities. This work was funded by the Bill & Melinda Gates Foundation (INV-043758), the National Institute of Allergy and Infectious Disease (U54AI170856, 1P01AI167966, U19AI181881), ARPA-H (P023), the Howard Hughes Medical Institute, the Audacious Project at the Institute for Protein Design, the Shurl and Kay Curci Foundation (H.E.E.), the Swedish Research Council (S.O.), and the Open Philanthropy Project Improving Protein Design Fund.

## Materials and Methods

### Docking

From the pdb file containing Crown_C5_-1, which had its five-fold rotational symmetry axis aligned to the z axis, we removed all chains but 3 chains of one ABC heterotrimer and truncated the C component by 1-4 helices. Because RPXdock is limited to single-chain inputs, for each instance of the heterotrimer with truncated C component, we connected amino acid residues of all three chains of the heterotrimers into one chain and saved as a new pdb. These pdb structures were used as input files for docking into D5_5 by restricting sampling to rotation and translation along the z axis using RPXdock^54^. The top scoring and the output with the highest shape compatibility was taken for the further design.

### Asymmetric interface design

The design of an asymmetrical C-D interface was carried out using the deep learning-based protein sequence design software ProteinMPNN (available at https://github.com/dauparas/ProteinMPNN). We used ProteinMPNN without biases, ProteinMPNN with adding biases to specific amino acids per residue position, and multistate ProteinMPNN. For each method, 100 sequences were designed across different temperatures (0.1, 0.2, 0.4, 0.5, 0.6, 0.8, and 1.0). In the ProteinMPNN with bias per residue position approach, we increased the likelihood of specific amino acids occupying predefined positions using a biasing script (available at https://github.com/dauparas/ProteinMPNN/blob/main/helper_scripts/make_bias_per_res_dict.py) with a small modification. This script normally favors certain amino acids at specific positions in one chain while disfavoring them in another chain. We modified it to apply positive biases for both chains to increase the probability of desired amino acids being incorporated on a predefined side. Specifically, we categorized the amino acids S, T, N, Q, V, I, L as small; F, Y, W as bulky; D, E as negatively charged; and R, H, K as positively charged. We biased amino acids in three ways: in “charges” approach we favored positively charged amino acids on one side and negatively charged on the other side, in “clashes” approach we favored small amino acids on one side and bulky on the other side, and in “charges-clashes” we favored positively charged and bulky amino acids on one side and negatively charged and small amino acids on the other side. Bias values of 0.1, 0.2, 0.69, 1.1, and 3.9 were used for both sides in each biasing approach, where 0.69 was for two-fold, 1.1 for three-fold and 3.9 for four-fold increase in likelihood to incorporate desired amino acid. Multi-state ProteinMPNN design was performed as previously described^39^ using beta values = −1, −0.5, −0.25.

### Extension of the C component using RFdiffusion

Translation of the portion of (ABC)_5_ cyclic assemblies was done in PyMOL Molecular Graphics System (Version 2.5.8, Schrödinger, LLC). For the full structure extension process, we reasoned that extending only one C component is sufficient, given that C components are organized in C5 symmetry and there are no neighboring subunits with which extensions might clash. For easier computing and to streamline the design filtration process, only four helices of the translated portion of the C component and two isolated helices belonging to the C component were used as an input for RFdiffusion. For each extension distance value, we generated 100 backbones using RFdiffusion. The ranges of amino acids provided for diffusion to generate backbones within the gaps were as follows: 50-150, 100-250, 150-350, and 250-400 residues for extensions of 25, 50, 75, and 100 Å, respectively. For the 50 Å extension with a subsequent 25° rotation we used a range of 150-350 residues. The range of amino acids needed to fill the gaps was initially estimated visually for the 25 Å extensions, with an allowance of ±50 residues. For subsequent extensions, the range was determined based on AF2 predictions for the successful 25 Å extensions.

### Reorienting N termini using RFdiffusion

To reorient the N termini, we used RFdiffusion with block-adjacency^47^. Briefly, we started by modifying the PDB structure of ABC heterotrimer to mimic the desired structure with the reoriented N-terminus. This involved deleting three loops and adding new loops and one helix of the desired length in specific locations, which we treated as separate chains. These modifications were made using PyMOL. The manipulated PDB structure was then used to create an adjacency matrix, which provided information on which helices the newly built helix (using RFdiffusion) should interact with. RFdiffusion was subsequently used to rebuild the new loops and the helix, generating 100 new heterotrimer backbones.

### Construction of synthetic genes

All synthetic genes were purchased from GenScript Inc. (Piscataway, NJ, USA). They were codon optimized for expression in E. coli and cloned into pET29b+ plasmid between Ndel/Xhol sites. Only A component had 6xHis-tag on C-terminus for facilitating IMAC purification of the complexes. C and D components had mScarlet and mNeonGreen fused to the N-terminus, respectively, for easier detection by SEC and SDS-PAGE gel.

### Protein expression

100 ng of each plasmid obtained from GenScript was diluted in 20 µL of DNase free water (Cytiva). 0.5 µL of each plasmid was used for the transformation of 5 µL of BL21(DE3)Star E. coli expression strain (Invitrogen) according to the manufacturer’s protocol. One bacterial colony was inoculated in 5 mL of Luria-Bertani (LB) medium containing 100 µg/mL kanamycin and grown overnight at 37°C with shaking at 225 rpm. 1 mL of the overnight culture was transferred to 50 mL Terrific Broth II media (MP Biomedical Cat# MP113046052) supplemented with 100 µg/mL kanamycin in 250 mL flasks. Bacteria was grown at 37°C until the optical density was 0.6-0.8 OD and then protein expression was induced by adding IPTG, after which the temperature was decreased to 18°C. Expression was going for 20-24 hours with shaking at 225 rpm.

### Immobilized metal affinity chromatography (IMAC)

50 mL cultures were harvested by centrifugation at 4000 *g* for 20 min at 4°C. Palleted bacteria were resuspended in 10 mL of lysis buffer (50 mM Tris pH 8.0, 300 mM NaCl, 20 mM imidazole, 10% glycerol). Resuspensions of A, B and C or A, B and D components were mixed and 300 µL of PMSF (100 mM in 20 % ethanol) was added to mixtures. Immediately upon adding PMSF, mixtures were sonicated at 65 % power for 5 minutes, with 10 seconds of on/off pulse. Lysed bacteria were left on ice for 30 minutes to allow formation of ABC heterotrimers and (ABC)_5_ or (ABD)_5_ cyclic assemblies, and subsequently centrifuged at 14000 *g* for 30 minutes at 18 °C. Following centrifugation, supernatant was kept and an additional 30 minutes left on room temperature to ensure proper assembly of (ABC)_5_ or (ABD)_5_ cyclic assemblies. Subsequently, supernatant was applied to 1 mL of Ni-NTA resin (Qiagen) for gravity chromatography, which was pre-equilibrated with 5 mL of lysis buffer. Columns were washed with 15 mL of wash buffer (50 mM Tris pH 8.0, 300 mM NaCl, 40 mM imidazole, 10% glycerol) and the protein was eluted with 1.7 mL of elution buffer (50 mM Tris pH 8.0, 300 mM NaCl, 300 mM imidazole, 300 mM EDTA, 10% glycerol). Only the last 1.3 mL of the elution was collected for further purification.

### Size-exclusion chromatography (SEC)

1.3 mL of sample obtained by IMAC purification was further purified using a Superose 6 10/300 Increase column (Cytiva) in SEC buffer (50 mM Tris pH 8.0, 300 mM NaCl) using ÄKTA Pure chromatography system. In addition to 280 nm absorbance, absorbance at 506 nm and 569 nm was followed. Upon mixing (ABC)_5_ and (ABD)_5_ cyclic assemblies, a second SEC purification was performed using the same buffer.

### SDS-PAGE

Peak fractions of cyclic assemblies and D5 assemblies were analyzed by SDS-PAGE electrophoresis. 1-5 µg of target protein was resuspended in 2× Laemmli Sample Buffer (Bio-Rad) with adding 1:20 Beta-mercaptoethanol and 15 µL of the mixture was added onto Any kD™ Criterion™ TGX Stain-Free™ Protein gel (Bio-Rad). 5 µL of Precision Plus Protein™ Unstained Protein Standards (Bio-Rad) was used and gels were run at 150 V for ∼50 min. Subsequently, gels were stained with GelCode Blue™ (Thermo Fisher Scientific) and destained in water. Stained gels were imaged using a Chemidoc XRS+ (Bio-Rad).

### Negative stain electron microscopy (nsEM)

3 µL of SEC purified samples of approximate concentration 0.05 mg/mL was deposited on 10 nm thick carbon film-coated 400 mesh copper grids (Electron Microscopy Sciences, CF400-Cu-TH) that was previously glow discharged for 20 seconds. Subsequently, grids were stained three times using 2% uranyl formate. Grids were screened using 120 kV Talos L120C transmission electron microscope. For collecting large datasets for obtaining 2D class averages and 3D reconstruction, E. Pluribus Unum (EPU) (FEI Thermo Scientific) software was used. Data processing was done using CryoSPARC™ versions v4.2.2, 4.4.0, and 4.4.1 (Structura Biotechnology Inc). We initially used D5 symmetry to generate the nsEM models, which then served as the starting point for the subsequent C1 reconstructions. Fitting of design models into corresponding density maps was done using ChimeraX ^63–65^.

### Dynamic light scattering (DLS)

To determine the size and uniformity of the particles, DLS measurements were performed using the Sizing and Polydispersity method on the Uncle instrument (Unchained Labs). 8.8 µL of SEC peak fractions were loaded into the provided glass cuvettes. DLS measurements were measured in triplicate at 25 °C, 10 acquisitions with each measuring 10 seconds in length. To determine the stability of the nanoparticles, SLS and intrinsic tryptophan fluorescence (ITF) (presented as the barycentric mean (BCM) of the emission spectrum) were measured in triplicate at 25 °C, followed by a thermal ramp from 25 °C to 95 °C at a ramp rate of 1.0 °C/min. Protein concentration (ranging from 0.1-0.4 mg/mL) and buffer conditions were accounted for in the software. Data was processed using Uncle Analysis Software version 6.01.0.0.

### Mass photometry (MP)

All mass photometry measurements were carried out on a TwoMP Auto mass photometer using the AcquireMP software (Refeyn). Proteins were diluted to ∼20 nM at least an hour before measurement in a flat-bottom 96-well polypropylene plate (Greiner). After centring the laser over a well on a 24-well gasket on commercially pre-cleaned slides (Refeyn), 5 µL of buffer (50 mM Tris, 300 mM NaCl, pH 8) was deposited into the well using the automated fluid handling system and used for finding focus using the drop dilution method. After focus was found, 5 µL of sample was pipetted into the gasket well and mixed once. 1-minute movies were recorded using the Normal field of view. Ratiometric contrast values for individual particles in each movie were measured and processed into mass distributions with DiscoverMP using a sample of 20 nM Beta-amylase (containing monomers (56 kDa), dimers (112 kDa), and tetramers (224 kDa) as a mass standards. Discover MP was used to fit gaussian distributions to the experimental mass distributions to calculate the mean mass of the particles.

### Cryo-EM sample preparation

To prepare the sample for Cryo-EM, 3 µL of the construct at a concentration of 1.5 mg/mL was pipetted onto a glow discharged 400 mesh copper ultrathin lacey carbon grid (Electron Microscopy Sciences, LC400-Cu-CC-25). The grid was immediately vitrified by plunge freezing into liquid ethane using a FEI Vitrobot Mk. IV at 22°C, 100% humidity, with a 7.5 second wait time and a 0.5 second blot time at −1 blot force. Subsequently, the grid was clipped and remained continuously submerged in liquid nitrogen until being loaded onto the microscope.

### Cryo-EM data processing

Using a 300 kV FEI Titan Krios equipt with a Gatan K3 direct electron detector and an Gatan BioQuantum energy filter, 4,871 movies were collected in SerialEM, utilizing beam shifts to collect 11 movies per stage movement at 105,000 times magnification with a pixel size of 0.843 Å/pixel. Image stacks were composed of 79 frames at an exposure rate of 0.0505 seconds/frame with a dose weight of 11.31 e^-^/A^2^/sec and an exposure time of 3.997 seconds resulting in a total dose of 45.21 e^−^/Å^2^. All movies were then imported into CryoSPARC v4.4 where all data processing took place. Firstly, exposures were preprocessed using patch motion correction, patch contrast transfer function estimation, and movies were curated, eliminating movies with CTF fit resolutions below 6 Å and those with average intensities above 472.77. Because of the central cavity and variable diameter of our proteins’ minor axis at different view angles, Blob Picking was unsuccessful. 177 particles were first manually selected, extracted to 800 pixels (952.4 Å) and classified into 2D class averages to generate templates. Two sequential rounds of template picking, extraction to 800 pixels, and 2D classification into 150 classes followed, resulting in a final population of 209,004 particles. Those particles were homogeneously refined in C5 symmetry using an ab-initio volume map of this construct previously characterized by nsEM which was low-pass filtered to 50 Å. Particles were downsampled to 400 pixels to reduce processing bandwidth before the volume was subsequently refined using Non-Uniform Refinement. Local CTF refinement was then performed on the particles followed by another round of Non-Uniform Refinement. To finalize the model, one last homogenous refinement was performed (generating our final half maps and global resolution of 4.30Å) which was then sharpened using a B-Factor of 239.90 (derived from our homogenous refinement) which was sharpened using DeepEMhancer ^66^ to generate our deposited coulombic potential map.

### Cryo-EM model building

To build our model, the computationally designed model was rigid-body docked into the final cryo-EM map in ChimeraX, and was subsequently trimmed to PolyA in Phenix^67,68^ and the backbone was relaxed into our volume map using Namdinator^69^. We then manually refined the backbone using ISOLDE^70^ in ChimeraX, and Coot^71^. Sidechains were trimmed to PolyA and a wwPDB validation service^72^ report was generated to verify that the model’s clashscore and Ramachandran outliers were each zero (**Supplementary Table 2**). The final structure was deposited in the Protein Data Bank (PDB) and Electron Microscopy Data Bank (EMDB) under accession numbers 9DZE and EMD-47327.

### Biolayer Interferometry (BLI)

BLI was performed on an Octet R8 (Sartorius). All biosensors were hydrated in a kinetics buffer (10 mM HEPES pH 7.4, 1% w/v Bovine Serum Albumin (BSA)). Biotinylated human IL-2Rβ (Acro Biosystems, ILB-H82E3) and human 4-1BB (Sino Biological, 10041-H27H-B) were diluted to a concentration of 2.5 μg/mL in the kinetics buffer and loaded onto streptavidin-coated biosensors (SAForteBio). Complexes were diluted in the kinetics buffer to a concentration of 15 nM and its association was measured for 300 seconds, followed by a dissociation for 300 seconds in the kinetics buffer. Data was processed using ForteBio Data Analysis Software version 9.0.0.10.

### Flow cytometry

200 μL of Streptavidin coated fluorescent Nile Red and Purple particles of nominal size 5.0-7.9 µm (Spherotech) were washed three times in phosphate buffered saline (PBS) containing 0.01% Tween 20 and 0.05% BSA according to the manufacturer protocol with spinning for 30 seconds at 21,000 x g. After washing, both particle types were resuspended in 250 μL of PBS with 0.01% Tween 20 and 0.05% BSA. Biotinylated IL-2Rβ and 4-1BB were incubated with Nile Red and Purple particles, respectively, at a concentration of 10 μg of receptor per 1 mg of Spherotech streptavidin particles, with gentle rotation for 30 minutes. The particles coated with biotinylated receptors were separated from unbound receptors by centrifugation at 21,000 x g for 30 seconds, followed by five washes in PBS with 0.01% Tween 20 and 0.05% BSA. Upon washing, particles were resuspended in 250 μL PBS with 0.01% Tween 20 and 0.05% BSA. Finally, 1 μL of each particle suspension was further resuspended in 18 μL of PBS with 0.01% Tween 20 and 0.05% BSA, to which 100 μL of the sample was added. The mixture was incubated at room temperature for 30 min and colocalization events were acquired using an AttuneNXT flow cytometer (ThermoFisher). Data files were analyzed using FlowJo software version 10 (BD Biosciences). Statistical analysis was performed using one-way ANOVA with Geisser-Greenhouse correction followed by Tukey’s multiple comparisons test, with individual variances computed for each comparison.

### Fluorescence microscopy

Coating streptavidin-coated fluorescent Nile Red and Purple particles with biotinylated human IL-2Rβ (Acro Biosystems, ILB-H82E3) and human 4-1BB (Sino Biological, 10041-H27H-B) was performed following the same procedure as for the flow cytometry experiments. 1 μL of each particle suspension was resuspended in 18 μL of PBS containing 0.01% Tween 20 and 0.05% BSA. Then, 4 μL of this mixture was incubated with 20 μL of the sample at room temperature for 30 minutes. Subsequently, the mixture was placed in a glass-bottomed 18-well Ibidi wellslide (Ibidi) and the particles were allowed to settle for 10 minutes prior to imaging. Images were acquired on an IN Cell Analyzer 2500HS microscope using a Nikon Plan Apo, CFI/60 20x objective, N.A. 0.75. Fluorescent particles were excited using LEDs emitting at 473 and 575 nm each with 10 msec exposure times. Emission was collected through bandpass filters centered at 525 (+/-24) nm and 623 (+/-12) nm, respectively. Each sample was imaged in 25 separate FOVs for quantification. Briefly, quantification was carried out using a custom python script which segments fluorescent particles in each channel and then, using one channel as a reference, measures the distance between each identified fluorescent particle and all fluorescent particles of the opposite color. Only fluorescent particles with a centroid-to-centroid separation distance of less than 7.5 µm (representing roughly twice the radius of the fluorescent particle) were kept as positive hits for fluorescent particle interaction. Percent fluorescent particle interactions were then determined by dividing the interacting fluorescent particles by the total fluorescent particle count for each channel in each image.

