## Supplements for "Computational design of bifaceted protein nanomaterials"

### Extended data

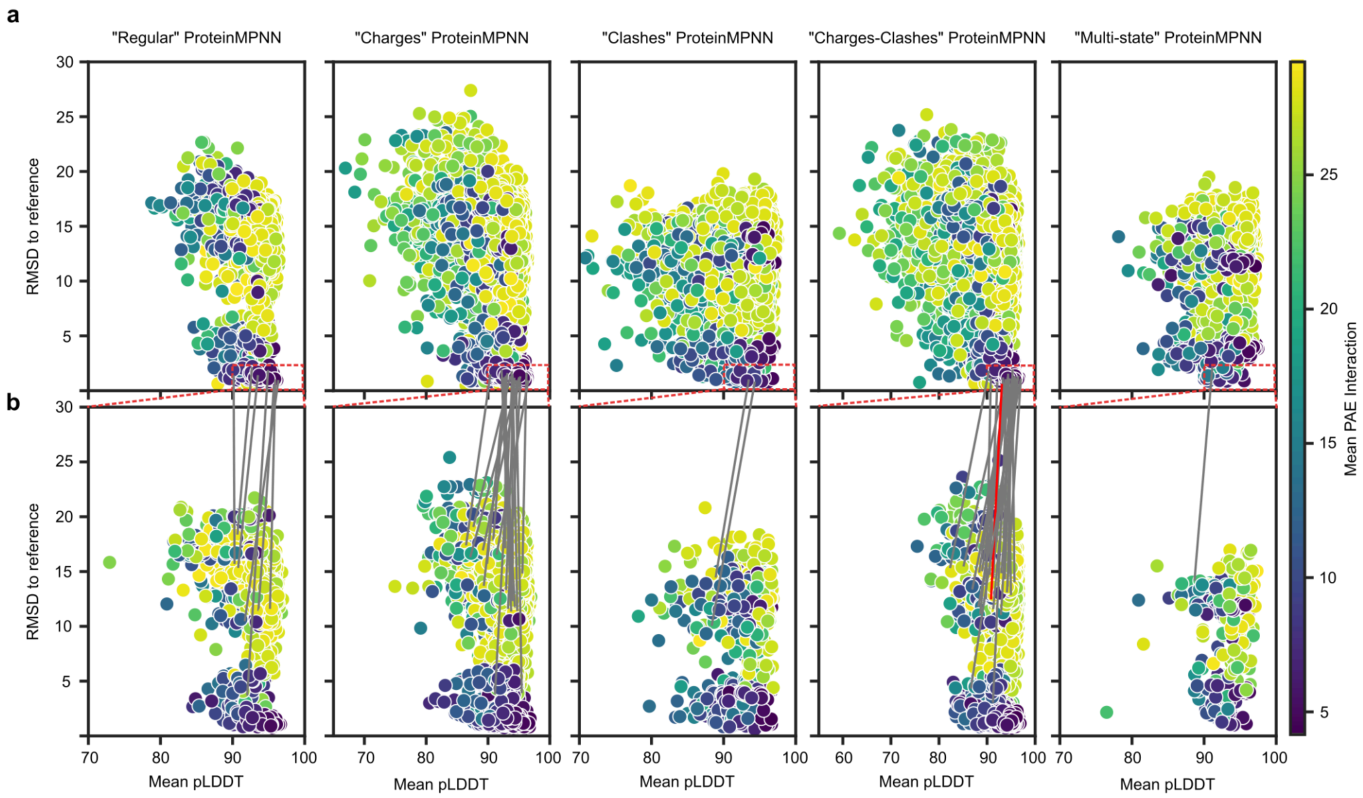


**Extended Data Fig. 1. AF2 structure prediction metrics of designed asymmetric C-D interface.**
**a,** AF2 metrics for the asymmetric C-D interface designs. The structures of the five C-terminal helices of the C and D subunits, comprising the designed interface, were predicted and pLDDT and pAE were plotted against RMSD to the computational design models (post-ProteinMPNN). Metrics for designs from each of the five ProteinMPNN strategies used to design asymmetric interfaces are shown separately. Each point represents a prediction from one of five AF2 models used to evaluate each design. Designs that passed the first round of filtering and were evaluated for off-target homotypic interface formation are boxed in red. **b,** AF2 metrics for off-target C-C or D-D interface formation using sequences that passed the first round of filtering. Only data corresponding to the best (i.e., lowest mean pAE interaction) off-target prediction for each design are shown. Gray lines connect data points for designs that were predicted to form the on-target C-D interface (**a**) and predicted to not form off-target C-C or D-D interfaces (**b**). The red line connects the on-target and off-target data points for pD5-14, which was experimentally verified to form exclusively $(ABC)_5$-$(ABD)_5$ complexes.

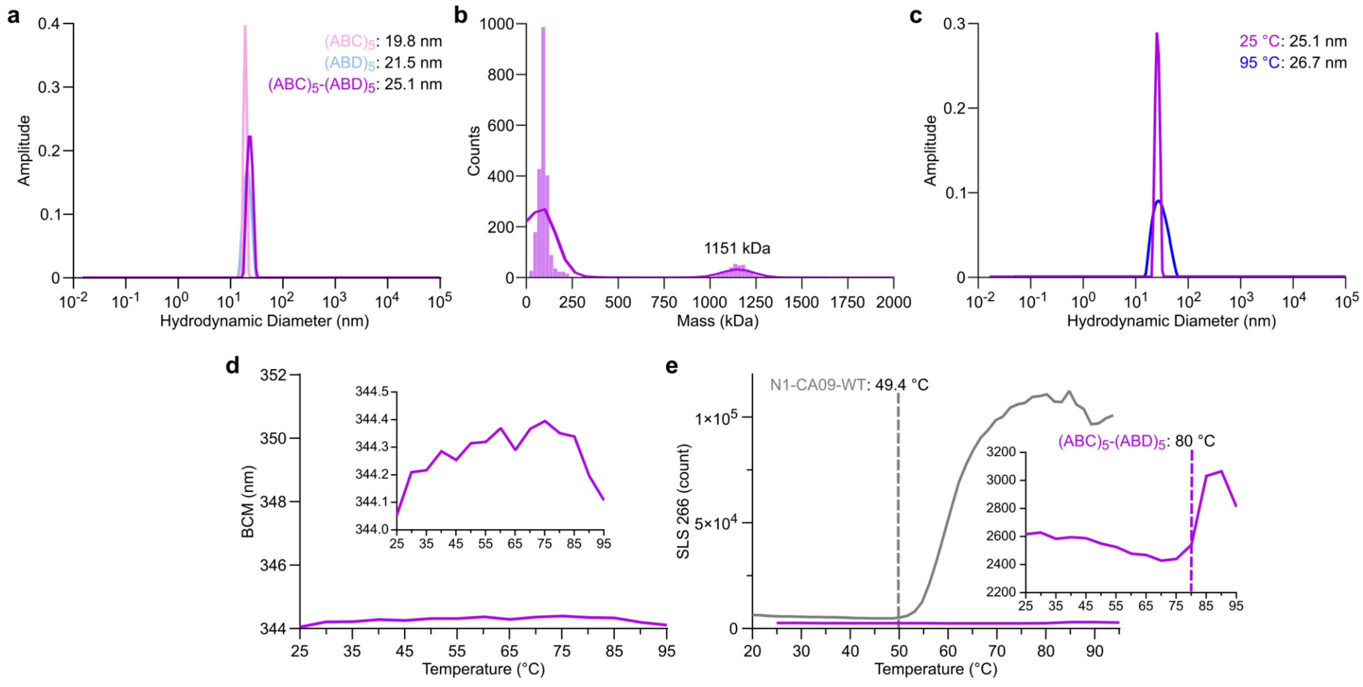


**Extended Data Fig. 2. Size distribution and thermal stability of pD5-14 bifaceted nanoparticles.** **a,** DLS and measured hydrodynamic diameters of pD5-14 $(ABC)_5$, $(ABD)_5$, and $(ABC)_5$-$(ABD)_5$. **b,** Mass photometry of pD5-14 $(ABC)_5$+$(ABD)_5$, showing an observed mass of 1151 kDa. The expected mass for the 30-subunit assembly is 1252 kDa. **c,** DLS measurements of pD5-14 $(ABC)_5$-$(ABD)_5$ at 25 °C and 95 °C. **d,** Nano differential scanning fluorimetry of $(ABC)_5$-$(ABD)_5$, plotted as the barycentric mean (BCM) of the emission spectrum during heating from 25–95 °C. The y axis spans the range of BCM values typically observed during protein denaturation, while the inset zooms in on the y axis to show the details of the data. **e,** Evaluation of thermal aggregation of $(ABC)_5$-$(ABD)_5$, measured by scattering intensity at 266 nm during heating from 25–95 °C. Thermal aggregation of an N1 influenza neuraminidase ectodomain (N1-CA09-WT)[73] is shown for comparison, while the inset zooms in on the y axis to show the details of the data $(ABC)_5$-$(ABD)_5$. Aggregation temperatures, as determined by UNcle Analysis Software, are indicated by dashed lines.

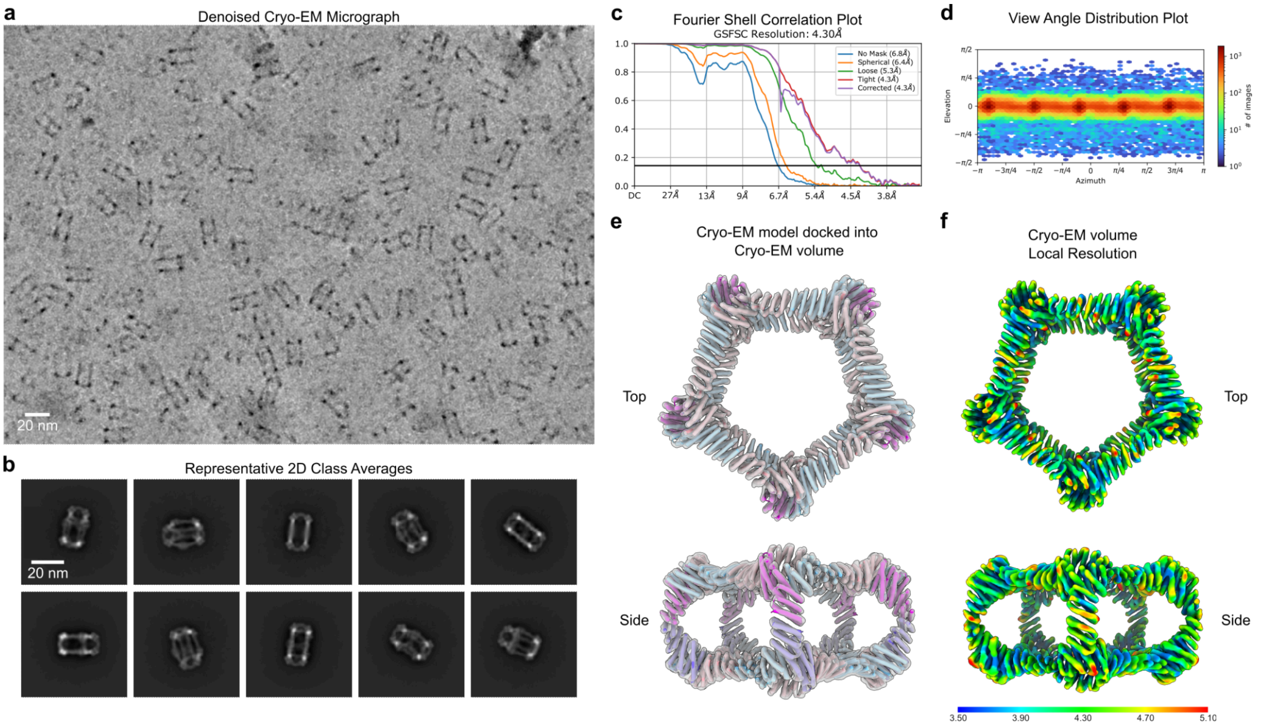


**Extended Data Fig. 3. Details of cryo-EM data processing.**
**a,** Denoised representative cryo-EM micrograph. **b,** Representative 2D class averages with scale bar showing particles from multiple view angles, exemplifying preference for ‘Side’ views over ‘Top’ views. **c,** Fourier shell correlation plot of volume map, illustrating 4.30 Å resolution estimation at FSC 0.143 cutoff. **d,** View angle distribution plot, demonstrating spatial preference for ‘Side’ views over ‘Top’ views. **e,** Cryo-EM model docked into cryo-EM volume. **f,** Local resolution estimation (Å) of cryo-EM volume. Scale bars: 20 nm.

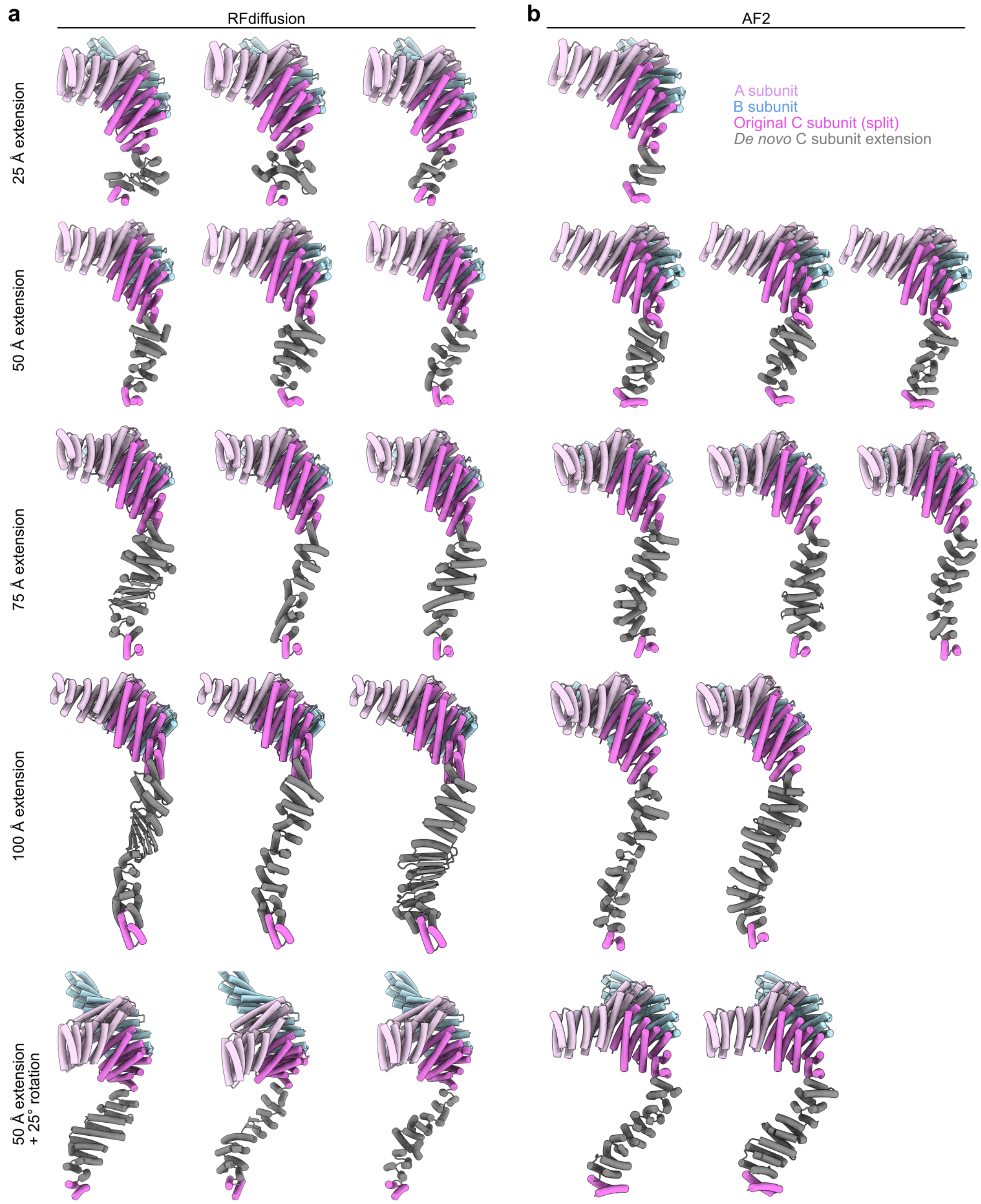


**Extended Data Fig. 4. Representative RFdiffusion and AF2 outputs of *de novo* extensions.**
**a,** Backbone structures of ABC heterotrimers output from RFdiffusion for different extension lengths. The images are oriented such that the two C-terminal helices of the C subunit that make up the pseudo-dihedral interface (pink) are at bottom. The *de novo* extensions that connect the C-terminal helices to the rest of the C subunit are colored gray. RFdiffusion generated diverse structures including both α-helices and β-sheets. **b,** Representative extended ABC heterotrimers that passed AF2 filtering. The *de novo* extensions of passing designs generally had well-packed α-helical repeat structures.

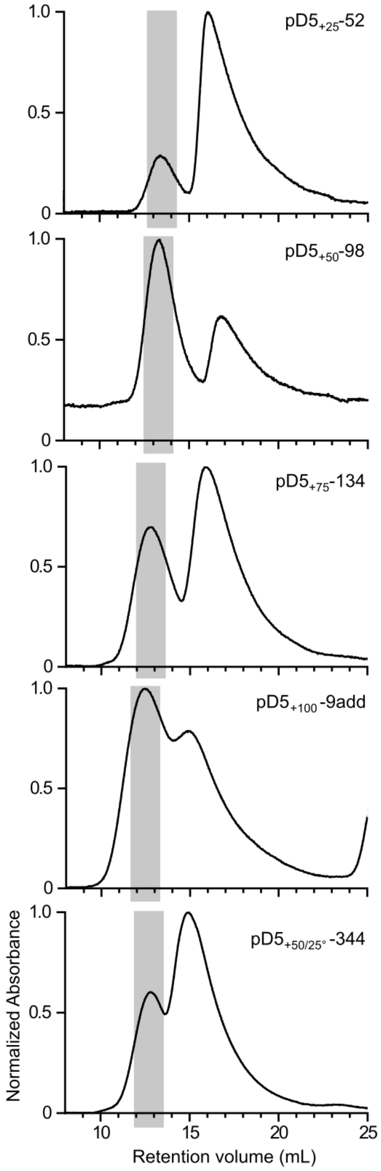


**Extended Data Fig. 5. SEC of extended bifaceted pD5 nanoparticles shown in Figure 4.**
Preparative SEC chromatograms are shown for each of the five extended pD5 nanoparticles shown in Figure 4. The gray rectangles depict the fractions pooled for further characterization.

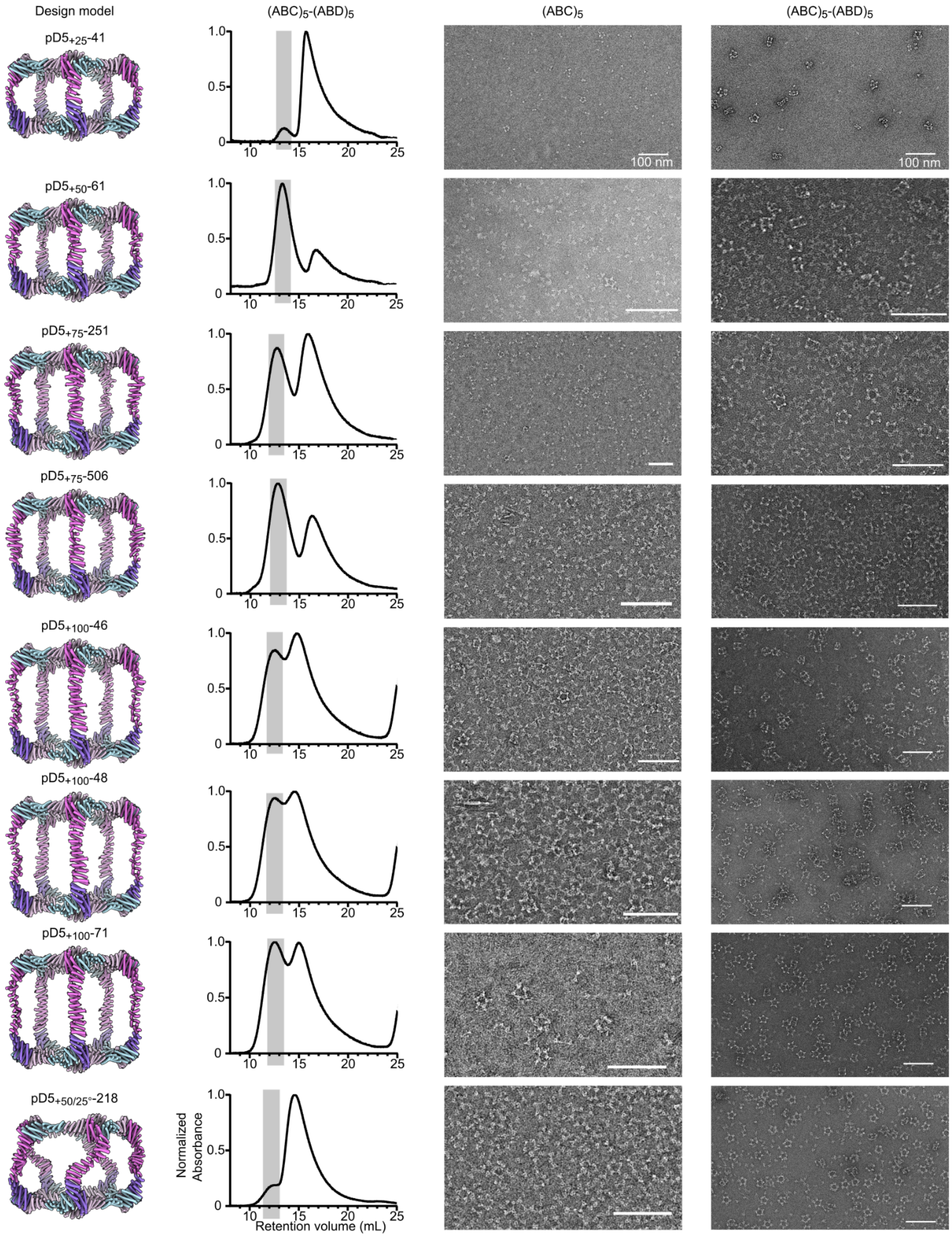


**Extended Data Fig. 6. Experimental characterization of additional extended bifaceted pD5 nanoparticles.**

*From top to bottom:* Additional assemblies extended by 25, 50, 75, and 100 Å, or extended by 50 Å and rotated by 25°. *From left to right:* computational design models, preparative SEC chromatograms, raw micrographs of $(ABC)_5$, and raw micrographs of $(ABC)_5$-$(ABD)_5$. Scale bars: 100 nm.

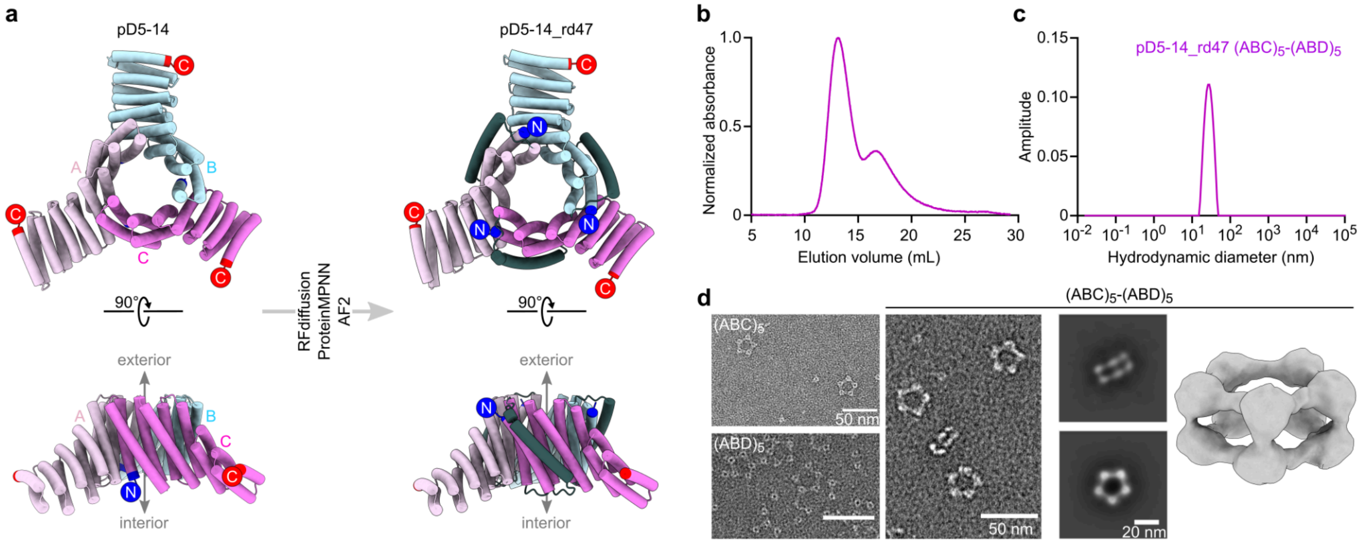


**Extended Data Fig. 7. Experimental characterization of pD5-14_rd47.**
**a,** Schematic of pD5-14 ABC heterotrimer redesign to generate pD5-14_rd47 ABC heterotrimers with exterior-facing N termini. The newly diffused α-helix is shown in dark gray, and N and C termini are indicated by blue and red circles, respectively. **b,** Preparative SEC chromatogram of pD5-14_rd47. **c,** DLS of SEC-purified pD5-14_rd47 $(ABC)_5$-$(ABD)_5$. **d,** nsEM characterization of pD5-14_rd47. *Left:* Raw micrographs are shown for the $(ABC)_5$ and $(ABD)_5$ components as well as the SEC-purified $(ABC)_5$-$(ABD)_5$ assemblies. *Right:* 2D class averages and a 3D reconstruction of pD5-14_rd47 $(ABC)_5$-$(ABD)_5$. Scale bars: 50 nm (raw micrographs) and 20 nm (2D class averages).

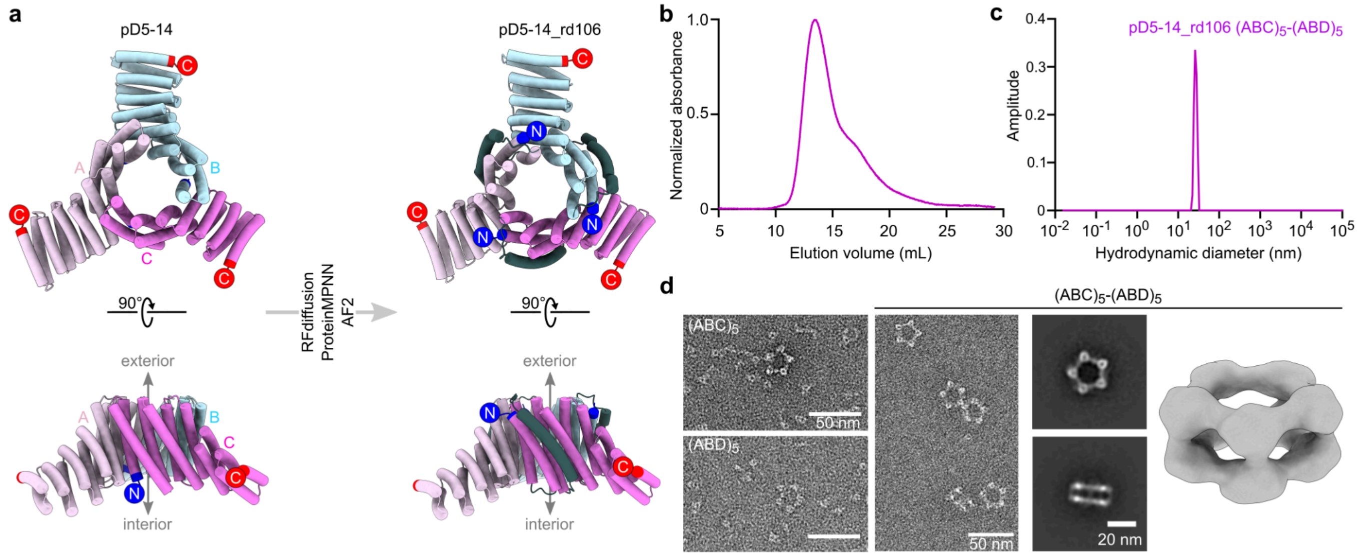


**Extended Data Fig. 8. Characterization of pD5-14_rd106.**

**a,** Schematic of pD5-14 ABC heterotrimer redesign to generate pD5-14_rd106 ABC heterotrimers with exterior-facing N termini. The newly diffused α-helix is shown in dark gray, and N and C termini are indicated by blue and red circles, respectively. **b,** Preparative SEC chromatogram of pD5-14_rd106. **c,** DLS of SEC-purified pD5-14_rd106 $(ABC)_5$-$(ABD)_5$. **d,** nsEM characterization of pD5-14_rd106. *Left:* Raw micrographs are shown for the $(ABC)_5$ and $(ABD)_5$ components as well as the SEC-purified $(ABC)_5$-$(ABD)_5$ assemblies. *Right:* 2D class averages and a 3D reconstruction of pD5-14_rd106 $(ABC)_5$-$(ABD)_5$. Scale bars: 50 nm (raw micrographs) and 20 nm (2D class averages).

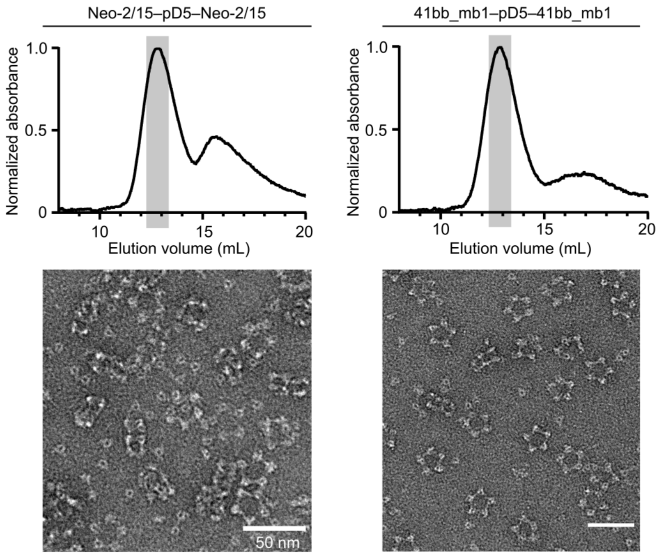


**Extended Data Fig. 9. SEC and nsEM of Neo-2/15–pD5–Neo-2/15 and 41bb_mb1–pD5–41bb_mb1 assemblies.**

*Top:* Preparative SEC chromatograms of Neo-2/15–pD5–Neo-2/15 and 41bb_mb1–pD5-14–41bb-mb1. The fractions collected for further characterization are marked by gray rectangles. *Bottom:* nsEM raw micrographs of the SEC-purified assemblies. Scale bar: 50 nm for both micrographs.

**Supplementary Table 1. Amino acid sequences of novel proteins used in this study.**

| ID | Subunit | Sequence |
|---|---|---|
| pD5-14 | A_6×His-tag | MGSLELALKALQILVNAAYVLAEIARDRGNEELLEKAARLAEEAARQAE RIARQARKEGNLELALKALQILVNAAYVLAEIARDRGNEELLEYAARLAE EAARQAIEIWAQAMEEGNQQLRTKAAHIILRAAEVLLEIARDRGNQELL EKAASLVDAVAALQQAAAAILEGDVEKAVRAAQEAVKAAKEAGDNDML RAVAIAALRIAKEAEKQGNVEVAVKAARVAVEAAKQAGDNDVLRKVAEQ ALRIAKEAEKQGNVEVAVKAARVAVEAAKQAGDNDVLRKVADQALEIAK AALEQGDIDVAQKAMDVAVEALTQAGGSGGSHHHHHH |
| | B | MGSPRLVLRALENMVRAAHTLAEIARDNGNEEWLERAARLAEEVA RRAERLAREARKEGNLELALKALQILVNAAYVLAEIARDRGNEEELE YAARLAEEAARQAIEIAAQAMEEGNLELALKALQIIVNAAYVLAEIAR DRGNEELLEKAASLAEAAAALAEAIAAILEGDVEKAVRAAQEAVKAA KEAGDNDMLRAVAIAALRIAKEAEKQGNVEVAVKAARVAVEAAKQA GDNDVLRKVAEQALRIAKEAEKQGNVEVAVKAARVAVEAAKQAGD NDVLRKVAEQALEIAKKAAEQGDVGVMQKAMDVALRAAGQAG |
| | mScarlet-I_C | MGSKGEAVIKEFMRFKVHMEGSMNGHEFEIEGEGEGRPYEGTQT AKLKVTKGGPLPFSWDILSPQFMYGSRAFIKHPADIPDYYKQSFPE GFKWERVMNFEDGGAVTVTQDTSLEDGTLIYKVKLRGTNFPPDGP VMQKKTMGWEASTERLYPEDGVLKGDIKMALRLKDGGRYLADFKT TYKAKKPVQMPGAYNVDRKLDITSHNEDYTVVEQYERSEGRHSTG GMDELYKGSGSGPELFLQDLRSLVEAARILARLARQRGDEHALER AARWAEQAARQAERLARQARKEGNLELALKALQILVNAAYVLAEIA RDRGNEELLEYAARLAEEAARQAIEIAAQAMEEGNFELALEALEIIN EAARVLARIAHHRGNQELLEKAASLTHASAALSRAIAAILEGDVEKA VRAAQEAVKAAKEAGDNDMLRAVAIAALRIAKEAEKQGNVEVAVKA ARVAVEAAKQAGDNDVLRLVSERALSIAASSVKQGNYEVKEKAIRV AKEANKQAG |
| | mNeonGreen_D | MGSKGEEDNMASLPATHELHIFGSINGVDFDMVGQGTGNPNDGY EELNLKSTKGDLQFSPWILVPHIGYGFHQYLPYPDGMSPFQAAMV DGSGYQVHRTMQFEDGASLTVNYRYTYEGSHIKGEAQVKGTGFPA DGPVMTNSLTAADWCRSKKTYPNDKTIISTFKWSYTTGNGKRYRS TARTTYTFAKPMAANYLKNQPMYVFRKTELKHSKTELNFKEWQKA FTDVMGMDELYKGSGSGPELFLQDLRSLVEAARILARLARQRGDE HALERAARWAEQAARQAERLARQARKEGNLELALKALQILVNAAYV LAEIARDRGNEELLEYAARLAEEAARQAIEIAAQAMEEGNFELALEA LEIINEAARVLARIAHHRGNQELLEKAASLTHASAALSRAIAAILEGD VEKAVRAAQEAVKAAKEAGDNDMLRAVAIAALRIAKEAEKQGNVEV AVKAARVAVEAAKQAGDNDVLKRVSETLLSIAAEATKQGNSEVMEK AIRVSEEAEKQAG |
| pD5$_{+25}$-52 | mScarlet-I_C | MGSKGEAVIKEFMRFKVHMEGSMNGHEFEIEGEGEGRPYEGTQT AKLKVTKGGPLPFSWDILSPQFMYGSRAFIKHPADIPDYYKQSFPE GFKWERVMNFEDGGAVTVTQDTSLEDGTLIYKVKLRGTNFPPDGP VMQKKTMGWEASTERLYPEDGVLKGDIKMALRLKDGGRYLADFKT TYKAKKPVQMPGAYNVDRKLDITSHNEDYTVVEQYERSEGRHSTG GMDELYKGSGSGPELFLQDLRSLVEAARILARLARQRGDEHALER AARWAEQAARQAERLARQARKEGNLELALKALQILVNAAYVLAEIA RDRGNEELLEYAARLAEEAARQAIEIAAQAMEEGNFELALEALEIIN EAARVLARIAHHRGNQELLEKAASLTHASAALSRAIAAILEGDVEKA VRAAQEAVKAAKEAGDNDMLRAVAIAALRIAKEAEKQGNVEVAVKA ARVAVEAAKQAGDRELKAKGLWREARGHLELGNTDAFAESLAAML EAVGEEERAALARELRELAVRLLEEGGSLADLRALLAALAALGDND |

| | | |
|---|---|---|
| | | VLRLVSERALSIAASSVKQGNYEVKEKAIRVAKEANKQAG |
| pD5$_{+25}$-41 | mScarlet-I_C | MGSKGEAVIKEFMRFKVHMEGSMNGHEFEIEGEGEGRPYEGTQT AKLKVTKGGPLPFSWDILSPQFMYGSRAFIKHPADIPDYYKQSFPE GFKWERVMNFEDGGAVTVTQDTSLEDGTLIYKVKLRGTNFPPDGP VMQKKTMGWEASTERLYPEDGVLKGDIKMALRLKDGGRYLADFKT TYKAKKPVQMPGAYNVDRKLDITSHNEDYTVVEQYERSEGRHSTG GMDELYKGSGSGPELFLQDLRSLVEAARILARLARQRGDEHALER AARWAEQAARQAERLARQARKEGNLELALKALQILVNAAYVLAEIA RDRGNEELLEYAARLAEEAARQAIEIAAQAMEEGNFELALEALEIIN EAARVLARIAHHRGNQELLEKAASLTHASAALSRAIAAILEGDVEKA VRAAQEAVKAAKEAGDNDMLRAVAIAALRIAKEAEKQGNVEVAVKA ARVAVEAAKQAGDVDLREEGREGQARGLIKLGDREGARKVLKEISS DAAEELLLELVTQRPLDVDLLRALMKHVEDNDVLRLVSERALSIAAS SVKQGNYEVKEKAIRVAKEANKQAG |
| pD5$_{+50}$-98 | mScarlet-I_C | MGSKGEAVIKEFMRFKVHMEGSMNGHEFEIEGEGEGRPYEGTQT AKLKVTKGGPLPFSWDILSPQFMYGSRAFIKHPADIPDYYKQSFPE GFKWERVMNFEDGGAVTVTQDTSLEDGTLIYKVKLRGTNFPPDGP VMQKKTMGWEASTERLYPEDGVLKGDIKMALRLKDGGRYLADFKT TYKAKKPVQMPGAYNVDRKLDITSHNEDYTVVEQYERSEGRHSTG GMDELYKGSGSGPELFLQDLRSLVEAARILARLARQRGDEHALER AARWAEQAARQAERLARQARKEGNLELALKALQILVNAAYVLAEIA RDRGNEELLEYAARLAEEAARQAIEIAAQAMEEGNFELALEALEIIN EAARVLARIAHHRGNQELLEKAASLTHASAALSRAIAAILEGDVEKA VRAAQEAVKAAKEAGDNDMLRAVAIAALRIAKEAEKQGNVEVAVKA ARVAVEAAKQAGNGALEAEAMKAELRAAARCMAEKGWSIEELEEL LKEVEKLGTAARAECYKEAALELARAIIERPEDEEAVELLREVLDKIIE LDASILNEILHELARAAIENEKHRAEMAAMAARVLRELEEVEGRDEL RERLLELLLEESLEEEALRELARCWVLLARDDEEGFREALERLRSL PLDAQVRRLRALAEAAEEQGDNDVLRLVSERALSIAASSVKQGNYE VKEKAIRVAKEANKQAG |
| pD5$_{+50}$-61 | mScarlet-I_C | MGSKGEAVIKEFMRFKVHMEGSMNGHEFEIEGEGEGRPYEGTQT AKLKVTKGGPLPFSWDILSPQFMYGSRAFIKHPADIPDYYKQSFPE GFKWERVMNFEDGGAVTVTQDTSLEDGTLIYKVKLRGTNFPPDGP VMQKKTMGWEASTERLYPEDGVLKGDIKMALRLKDGGRYLADFKT TYKAKKPVQMPGAYNVDRKLDITSHNEDYTVVEQYERSEGRHSTG GMDELYKGSGSGPELFLQDLRSLVEAARILARLARQRGDEHALER AARWAEQAARQAERLARQARKEGNLELALKALQILVNAAYVLAEIA RDRGNEELLEYAARLAEEAARQAIEIAAQAMEEGNFELALEALEIIN EAARVLARIAHHRGNQELLEKAASLTHASAALSRAIAAILEGDVEKA VRAAQEAVKAAKEAGDNDMLRAVAIAALRIAKEAEKQGNVEVAVKA ARVAVEAAKQAGDEELYQRALAMELASLLKRGDYEEAKELIEREPIT EEATRIMCEILKNDLKALHRAAKLLLEAGLREGAKLFAKSMVEGVKK GLGSKEGVKCLNELADELEFSQEECDLLADLLVEAGEEAYEEGDEE ELEETVKLLAEWLEKCQISAATARRILEWMERLSLEQQVELLAAMV KSQTDNDVLRLVSERALSIAASSVKQGNYEVKEKAIRVAKEANKQA G |
| pD5$_{+75}$-134 | mScarlet-I_C | MGSKGEAVIKEFMRFKVHMEGSMNGHEFEIEGEGEGRPYEGTQT AKLKVTKGGPLPFSWDILSPQFMYGSRAFIKHPADIPDYYKQSFPE GFKWERVMNFEDGGAVTVTQDTSLEDGTLIYKVKLRGTNFPPDGP VMQKKTMGWEASTERLYPEDGVLKGDIKMALRLKDGGRYLADFKT TYKAKKPVQMPGAYNVDRKLDITSHNEDYTVVEQYERSEGRHSTG GMDELYKGSGSGPELFLQDLRSLVEAARILARLARQRGDEHALER AARWAEQAARQAERLARQARKEGNLELALKALQILVNAAYVLAEIA RDRGNEELLEYAARLAEEAARQAIEIAAQAMEEGNFELALEALEIIN |

| | | |
|---|---|---|
| | | EAARVLARIAHHRGNQELLEKAASLTHASAALSRAIAAILEGDVEKA<br>VRAAQEAVKAAKEAGDNDMLRAVAIAALRIAKEAEKQGNVEVAVKA<br>ARVAVEAAKQAGDDELTALGLAGQMKGLAKAGADLEELREVLDKLV<br>AKLEELEEVSEETKDRLTDVLEELLKREEYLELMREAMRRCKELGL<br>LEVLERLTERVIEEGASVEQLRELMEAYRELGLREEAQRLAVEAME<br>KLIEAGDLEGLLELLEEMLEAAEVFSRDLLIDLLLRAARVMLERMKE<br>ADLEERGEIAEQLVRTAEMLLEVVEDDAAALRELARLLEEAMRLLAE<br>LHTEDGIEWAGEQLAATAALAARALIALGDLEGFRRLMDELAEQLE<br>GDLELQVEVLQALRDAAADRDNDVLRLVSERALSIAASSVKQGNYE<br>VKEKAIRVAKEANKQAG |
| pD5$_{+75}$-251 | mScarlet-I_C | MGSKGEAVIKEFMRFKVHMEGSMNGHEFEIEGEGEGRPYEGTQT<br>AKLKVTKGGPLPFSWDILSPQFMYGSRAFIKHPADIPDYYKQSFPE<br>GFKWERVMNFEDGGAVTVTQDTSLEDGTLIYKVKLRGTNFPPDGP<br>VMQKKTMGWEASTERLYPEDGVLKGDIKMALRLKDGGRYLADFKT<br>TYKAKKPVQMPGAYNVDRKLDITSHNEDYTVVEQYERSEGRHSTG<br>GMDELYKGSGSGPELFLQDLRSLVEAARILARLARQRGDEHALER<br>AARWAEQAARQAERLARQARKEGNLELALKALQILVNAAYVLAEIA<br>RDRGNEELLEYAARLAEEAARQAIEIAAQAMEEGNFELALEALEIIN<br>EAARVLARIAHHRGNQELLEKAASLTHASAALSRAIAAILEGDVEKA<br>VRAAQEAVKAAKEAGDNDMLRAVAIAALRIAKEAEKQGNVEVAVKA<br>ARVAVEAAKQAGDIELLCQAATGLCRGAAKLMVEGKELTDVEEFRE<br>LAAELKELKEELNRLEVARALASLYALLYMTGDEEAREELLEATREM<br>REMVASDRAKRQELLDELSEQVEALDGCGIVDEELLLLLAEEALEL<br>GDLELFEEVLKRALETDEETLERFIEILEEAGDLEALLLAAEVLLEDA<br>LERGREGAVRCLARLLRRLAELGAGADSLLALLQRCLRLPEDFVVR<br>ILEEVVEELGEETLLRLAQQLLEQGLEEALVALAGGMARAGHAEALL<br>ELLQQLEEQGLEELSIRVLQAIVETAKDNDVLRLVSERALSIAASSVK<br>QGNYEVKEKAIRVAKEANKQAG |
| pD5$_{+75}$-506 | mScarlet-I_C | MGSKGEAVIKEFMRFKVHMEGSMNGHEFEIEGEGEGRPYEGTQT<br>AKLKVTKGGPLPFSWDILSPQFMYGSRAFIKHPADIPDYYKQSFPE<br>GFKWERVMNFEDGGAVTVTQDTSLEDGTLIYKVKLRGTNFPPDGP<br>VMQKKTMGWEASTERLYPEDGVLKGDIKMALRLKDGGRYLADFKT<br>TYKAKKPVQMPGAYNVDRKLDITSHNEDYTVVEQYERSEGRHSTG<br>GMDELYKGSGSGPELFLQDLRSLVEAARILARLARQRGDEHALER<br>AARWAEQAARQAERLARQARKEGNLELALKALQILVNAAYVLAEIA<br>RDRGNEELLEYAARLAEEAARQAIEIAAQAMEEGNFELALEALEIIN<br>EAARVLARIAHHRGNQELLEKAASLTHASAALSRAIAAILEGDVEKA<br>VRAAQEAVKAAKEAGDNDMLRAVAIAALRIAKEAEKQGNVEVAVKA<br>ARVAVEAAKQAGDEELYIEALAGEAKELVKLGDIEGAAKVFLEMVER<br>DLEKAAETAKEIIEEYQEEATELFLLLGKENLEALVEILKALKKMESM<br>TFEELLKAAKILGKVILEDESVSEEVCKIFIEFVKKILIQSEKATLEELL<br>EMAEVLQKILEKRELSEDIRREFVLCLSQVLLQIAKRIKELDVEEAEE<br>KIEELIEKLEKLQSEEELDEELMGQTCLILLETCITVLPIDDKRIEKISR<br>LVELLVSLSNIDIHRKACELLDAEIDKLVDAPLEYKLTMLACLRDVAAA<br>AGDNDVLRLVSERALSIAASSVKQGNYEVKEKAIRVAKEANKQAG |
| pD5$_{+100}$-9add | mScarlet-I_C | MGSKGEAVIKEFMRFKVHMEGSMNGHEFEIEGEGEGRPYEGTQT<br>AKLKVTKGGPLPFSWDILSPQFMYGSRAFIKHPADIPDYYKQSFPE<br>GFKWERVMNFEDGGAVTVTQDTSLEDGTLIYKVKLRGTNFPPDGP<br>VMQKKTMGWEASTERLYPEDGVLKGDIKMALRLKDGGRYLADFKT<br>TYKAKKPVQMPGAYNVDRKLDITSHNEDYTVVEQYERSEGRHSTG<br>GMDELYKGSGSGPELFLQDLRSLVEAARILARLARQRGDEHALER<br>AARWAEQAARQAERLARQARKEGNLELALKALQILVNAAYVLAEIA<br>RDRGNEELLEYAARLAEEAARQAIEIAAQAMEEGNFELALEALEIIN<br>EAARVLARIAHHRGNQELLEKAASLTHASAALSRAIAAILEGDVEKA<br>VRAAQEAVKAAKEAGDNDMLRAVAIAALRIAKEAEKQGNVEVAVKA |

| | | |
|---|---|---|
| | | ARVAVEAAKQAGDERLQSEALLAEAKALREMGDFDGFAETLRELLE<br>LPSAREVIKLEELLEFAQYLTQQRIAGADPELCDELLQELLEKFEEH<br>RLEMLTVIALELAKNPAFAERLKEVFRELVELGISEEKKKELLIYMFR<br>QGLVEEVLELCLEENVSEEILRELGLLAIDDPEVFEKLIDVMRTLGYT<br>RLARELLAQRLESLLVRPPLSPQQKQEMLDTAIRLMRECRELGEMS<br>PRERQLLFQCGQVFIHDLEAFQQLCTELQKLREAGIPEVNTLAFQLL<br>QYLLQNKDQLDVEDFAQQVEQLLLILDLEQFKEAVEVILQSLKSAPL<br>ELQVEVLEAIRRAAEKKGDNDVLRLVSERALSIAASSVKQGNYEVK<br>EKAIRVAKEANKQAG |
| pD5$_{+100}$-46 | mScarlet-I_C | MGSKGEAVIKEFMRFKVHMEGSMNGHEFEIEGEGEGRPYEGTQT<br>AKLKVTKGGPLPFSWDILSPQFMYGSRAFIKHPADIPDYYKQSFPE<br>GFKWERVMNFEDGGAVTVTQDTSLEDGTLIYKVKLRGTNFPPDGP<br>VMQKKTMGWEASTERLYPEDGVLKGDIKMALRLKDGGRYLADFKT<br>TYKAKKPVQMPGAYNVDRKLDITSHNEDYTVVEQYERSEGRHSTG<br>GMDELYKGSGSGPELFLQDLRSLVEAARILARLARQRGDEHALER<br>AARWAEQAARQAERLARQARKEGNLELALKALQILVNAAYVLAEIA<br>RDRGNEELLEYAARLAEEAARQAIEIAAQAMEEGNFELALEALEIIN<br>EAARVLARIAHHRGNQELLEKAASLTHASAALSRAIAAILEGDVEKA<br>VRAAQEAVKAAKEAGDNDMLRAVAIAALRIAKEAEKQGNVEVAVKA<br>ARVAVEAAKQAGDERLQAEALLAKAEALMEMGDFDGFAATLDELL<br>RLPEARAVVELERLKRFARHLTQQMIAGADPERCERLLQQLLEKFE<br>EHRLEMLTVIALELAKNPEFAEKLKEVFRELVELGISEEEKKELLLYM<br>FEQGLVELVLELCREENVSEEILEELAEKAIDNPEVFFKLIEVMRELG<br>YTELARKKLAERLKKLLEKPPLSPEEKKEMLDTAIELMEEARELGKL<br>TEEERELLYECMLVFVFDEEAFKKLCEELKKLREAGIEEVNELAYES<br>LLELLKRRDELDVDDFAKQVRDLLLILDEERFKEAVEEILKSLESAPL<br>ELQVKVLEALRKAAKEKGDNDVLRLVSERALSIAASSVKQGNYEVK<br>EKAIRVAKEANKQAG |
| pD5$_{+100}$-48 | mScarlet-I_C | MGSKGEAVIKEFMRFKVHMEGSMNGHEFEIEGEGEGRPYEGTQT<br>AKLKVTKGGPLPFSWDILSPQFMYGSRAFIKHPADIPDYYKQSFPE<br>GFKWERVMNFEDGGAVTVTQDTSLEDGTLIYKVKLRGTNFPPDGP<br>VMQKKTMGWEASTERLYPEDGVLKGDIKMALRLKDGGRYLADFKT<br>TYKAKKPVQMPGAYNVDRKLDITSHNEDYTVVEQYERSEGRHSTG<br>GMDELYKGSGSGPELFLQDLRSLVEAARILARLARQRGDEHALER<br>AARWAEQAARQAERLARQARKEGNLELALKALQILVNAAYVLAEIA<br>RDRGNEELLEYAARLAEEAARQAIEIAAQAMEEGNFELALEALEIIN<br>EAARVLARIAHHRGNQELLEKAASLTHASAALSRAIAAILEGDVEKA<br>VRAAQEAVKAAKEAGDNDMLRAVAIAALRIAKEAEKQGNVEVAVKA<br>ARVAVEAAKQAGDERLQAEALLGKAEALRKMGDFDGFAETLEELLT<br>LPSAVEVMKLEELLEFARYLTEQKIAGADPERCDRLLEELLEKFEEH<br>RLEMLKVIALELAKNPAFAEELKEVFRELVRLGMSEEEKKELLIYMF<br>EQGLVELVLELCLEEKVSEEILRELGEKAIDNPEVFFKLIDVMKELGY<br>IELARDMLAKKLESLLVKPPLSPEEKEELLKTAIELMDCAKELGYLTE<br>REKELLFECGEVFVFDQEAMEKLCEKLQELREAGIEEVNELAFELL<br>EYLLEHADELDVDDFAKQVEALLRILDKEQFKEAVEVIWESLESAPL<br>ERQVKILEAIRRAAVEAGDNDVLRLVSERALSIAASSVKQGNYEVKE<br>KAIRVAKEANKQAG |
| pD5$_{+100}$-71 | mScarlet-I_C | MGSKGEAVIKEFMRFKVHMEGSMNGHEFEIEGEGEGRPYEGTQT<br>AKLKVTKGGPLPFSWDILSPQFMYGSRAFIKHPADIPDYYKQSFPE<br>GFKWERVMNFEDGGAVTVTQDTSLEDGTLIYKVKLRGTNFPPDGP<br>VMQKKTMGWEASTERLYPEDGVLKGDIKMALRLKDGGRYLADFKT<br>TYKAKKPVQMPGAYNVDRKLDITSHNEDYTVVEQYERSEGRHSTG<br>GMDELYKGSGSGPELFLQDLRSLVEAARILARLARQRGDEHALER<br>AARWAEQAARQAERLARQARKEGNLELALKALQILVNAAYVLAEIA<br>RDRGNEELLEYAARLAEEAARQAIEIAAQAMEEGNFELALEALEIIN |

| | | |
|---|---|---|
| | | EAARVLARIAHHRGNQELLEKAASLTHASAALSRAIAAILEGDVEKAVRAAQEAVKAAKEAGDNDMLRAVAIAALRIAKEAEKQGNVEVAVKAARVAVEAAKQAGDEELEAEALLGKAEALMEMGDFEGFAETLRQLLELPSARRVMKLERLKRFARHLTQQMIAGADPELCEELLRQLLEKFREHRLECLEVIALELAKNPEFAEKWKEVFRELVELGISEEKRKELLIYAFEQGLVEEVLELCREERVSEEILRELGLLAIDDPEVFFRLIEVMRELGYLELAREMLAEKLRSLLEKPPLSPEEKEELLKTAIRLMECCRELGYMTPEERELLFECGQVFVFDEEAFKKLCEKLKELREAGIEEVNELAFELLQYLLEHRDELDVEELAEQIERLCLILNEEQFKEAVELILESLKSAPLELQVEVLEALRNAAKEKGDNDVLRLVSERALSIAASSVKQGNYEVKEKAIRVAKEANKQAG |
| pD5$_{+50/25°}$-344 | mScarlet-I_C | MGSKGEAVIKEFMRFKVHMEGSMNGHEFEIEGEGEGRPYEGTQTAKLKVTKGGPLPFSWDILSPQFMYGSRAFIKHPADIPDYYKQSFPEGFKWERVMNFEDGGAVTVTQDTSLEDGTLIYKVKLRGTNFPPDGPVMQKKTMGWEASTERLYPEDGVLKGDIKMALRLKDGGRYLADFKTTYKAKKPVQMPGAYNVDRKLDITSHNEDYTVVEQYERSEGRHSTGGMDELYKGSGSGPELFLQDLRSLVEAARILARLARQRGDEHALERAARWAEQAARQAERLARQARKEGNLELALKALQILVNAAYVLAEIARDRGNEELLEYAARLAEEAARQAIEIAAQAMEEGNFELALEALEIINEAARVLARIAHHRGNQELLEKAASLTHASAALSRAIAAILEGDVEKAVRAAQEAVKAAKEAGDNDMLRAVAIAALRIAKEAEKQGNVEVAVKAARVAVEAAKQAGDVALLAKGLVAKARSLVELGADLEEIKEVIREAIRAMVECKDEDCMRELAELMLELQKKGEKELFEFALREICRQLKDSDTEFRVRFLLICREVGLDVETLGRLTEELIRQGNNASELLSLLLERALERGDEDLADLVEEIAIKHRETAVAVARRLARLAEERDAATSARCRERLDRLAAHFADDEELGKALVEEAARRLRRLERIDEARQYYLETLERAARLGLKTIEELAALGTEVLMEKEEMKKEDVEKLLDALEEAEELKDSALVAALEALRKKAVEKGDNDVLRLVSERALSIAASSVKQGNYEVKEKAIRVAKEANKQAG |
| pD5$_{+50/25°}$-218 | mScarlet-I_C | MGSKGEAVIKEFMRFKVHMEGSMNGHEFEIEGEGEGRPYEGTQTAKLKVTKGGPLPFSWDILSPQFMYGSRAFIKHPADIPDYYKQSFPEGFKWERVMNFEDGGAVTVTQDTSLEDGTLIYKVKLRGTNFPPDGPVMQKKTMGWEASTERLYPEDGVLKGDIKMALRLKDGGRYLADFKTTYKAKKPVQMPGAYNVDRKLDITSHNEDYTVVEQYERSEGRHSTGGMDELYKGSGSGPELFLQDLRSLVEAARILARLARQRGDEHALERAARWAEQAARQAERLARQARKEGNLELALKALQILVNAAYVLAEIARDRGNEELLEYAARLAEEAARQAIEIAAQAMEEGNFELALEALEIINEAARVLARIAHHRGNQELLEKAASLTHASAALSRAIAAILEGDVEKAVRAAQEAVKAAKEAGDNDMLRAVAIAALRIAKEAEKQGNVEVAVKAARVAVEAAKQAGDVGLLALGLIAEARSLIELGAELEEIKEVIEEAIEAMVEDGRKESMKALAELAKYLQEKGEKELFEYLLEKLAEELKDKDTEFKVEVLLIFSEVGLDIETIEKLTRKLIEEGNNAAELLSELLARALERGDQELADLVREIAIEKRETVTAVAAELAARAQREDAATAARCEALLDELLAHFAEDEELGRTWVEQAAARLEELERIDDAIQFMLDTLERAAELGLRTIGDLAAKLMEVLMEKEEMKRADIEKLLDRLLEAESLRAEALVALLEAARDKAIEKGDNDVLRLVSERALSIAASSVKQGNYEVKEKAIRVAKEANKQAG |
| pD5-14_rd106 | A | MGSSELLRKAALLAAEAAEQAARIAKQAAKGELKNLELALKALQILVNAAYVLAEIARDRGEKPEIEEILPELRKLAKEAEEEAKEAKKEIEKATEQGLELALKALQILVNAAYVLAEIARDRGNEELLEIAAKLAEEAAELAIEVWALAMERGNQQLRTKAAHIILRAAEVLLEIARDRGNQELLEKAASLVDAVAALQQAAAAILEGDVEKAVRAAQEAVKAAKEAGDNDMLRAVAIAALRIAKEAEKQGNVEVAVKAARVAVEAAKQAGDNDVLRKVAEQALRIAKEAEKQGNVEVAVKAARVAVEAAKQAGDNDVLRKVADQALEIAKAALEQGDIDVAQKAMDVAVEALTQAGGSGGSHHHHHH |

| | | |
|---|---|---|
| | B | MGSEEWLTRAALLALEVAVRAARLAAEAAKGVRENPRLVLRALEN MVRAAHTLAEIARDNGPGTPEREEEIEPLIEELEKELERAKKAFEEY GKNPEGLELALKALQILVNAAYVLAEIARDRGNERLLEAAAKLAESA AELAIKVAAEAMELGNLELALKALQIIVNAAYVLAEIARDRGNEELLE KAASLAEAAAALAEAIAAILEGDVEKAVRAAQEAVKAAKEAGDNDM LRAVAIAALRIAKEAEKQGNVEVAVKAARVAVEAAKQAGDNDVLRKV AEQALRIAKEAEKQGNVEVAVKAARVAVEAAKQAGDNDVLRKVAE QALEIAKKAAEQGDVGVMQKAMDVALRAAGQAG |
| | mScarlet-I_C | MGSKGEAVIKEFMRFKVHMEGSMNGHEFEIEGEGEGRPYEGTQT AKLKVTKGGPLPFSWDILSPQFMYGSRAFIKHPADIPDYYKQSFPE GFKWERVMNFEDGGAVTVTQDTSLEDGTLIYKVKLRGTNFPPDGP VMQKKTMGWEASTERLYPEDGVLKGDIKMALRLKDGGRYLADFKT TYKAKKPVQMPGAYNVDRKLDITSHNEDYTVVEQYERSEGRHSTG GMDELYKGSGSGTHALVRAAKWAAQAAEQALRLAVQAAKGTVEN PELFLQDLRSLVEAARILARLARQRGEDTPEGDEELDRLAERLKEQ LRRLAEEFAAAAANSENLELALKALQILVNAAYVLAEIARDRGWERV LDAAAELAEAAARMAIEIAARAMEEGNFELALEALEIINEAARVLARI AHHRGNQELLEKAASLTHASAALSRAIAAILEGDVEKAVRAAQEAVK AAKEAGDNDMLRAVAIAALRIAKEAEKQGNVEVAVKAARVAVEAAK QAGDNDVLRLVSERALSIAASSVKQGNYEVKEKAIRVAKEANKQAG |
| | mNeonGreen_D | MGSKGEEDNMASLPATHELHIFGSINGVDFDMVGQGTGNPNDGY EELNLKSTKGDLQFSPWILVPHIGYGFHQYLPYPDGMSPFQAAMV DGSGYQVHRTMQFEDGASLTVNYRYTYEGSHIKGEAQVKGTGFPA DGPVMTNSLTAADWCRSKKTYPNDKTIISTFKWSYTTGNGKRYRS TARTTYTFAKPMAANYLKNQPMYVFRKTELKHSKTELNFKEWQKA FTDVMGMDELYKGSGSGTHALVRAAKWAAQAAEQALRLAVQAAK GTVENPELFLQDLRSLVEAARILARLARQRGEDTPEGDEELDRLAE RLKEQLRRLAEEFAAAAANSENLELALKALQILVNAAYVLAEIARDR GWERVLDAAAELAEAAARMAIEIAARAMEEGNFELALEALEIINEAA RVLARIAHHRGNQELLEKAASLTHASAALSRAIAAILEGDVEKAVRA AQEAVKAAKEAGDNDMLRAVAIAALRIAKEAEKQGNVEVAVKAARV AVEAAKQAGDNDVLKRVSETLLSIAAEATKQGNSEVMEKAIRVSEE AEKQAG |
| pD5-14_rd47 | A | MGSGELLFKAAKLALEAAKQAARIAAQAIKGEDNNLELALKALQILV NAAYVLAEIARDRGEKELTEEQKRELKELIEELKKELREALEVAKTEL GENLELALKALQILVNAAYVLAEIARDRGDEELLEAAAEIAESAAEIAI LVWAKAMEQGNQQLRTKAAHIILRAAEVLLEIARDRGNQELLEKAA SLVDAVAALQQAAAAILEGDVEKAVRAAQEAVKAAKEAGDNDMLRA VAIAALRIAKEAEKQGNVEVAVKAARVAVEAAKQAGDNDVLRKVAE QALRIAKEAEKQGNVEVAVKAARVAVEAAKQAGDNDVLRKVADQAL EIAKAALEQGDIDVAQKAMDVAVEALTQAGGSGGSHHHHHH |
| | B | MGSAEWLERAAELALEVAERAARLAAEAWRTGVEEPRLVLRALEN MVRAAHTLAEIARDNGERPDLEEIRERLEELAERLREELERAKRVA KEEEGKNLELALKALQILVNAAYVLAEIARDRGIELLLEAAAELAETA AELAIIAAAKAMEQGNLELALKALQIIVNAAYVLAEIARDRGNEELLE KAASLAEAAAALAEAIAAILEGDVEKAVRAAQEAVKAAKEAGDNDM LRAVAIAALRIAKEAEKQGNVEVAVKAARVAVEAAKQAGDNDVLRKV AEQALRIAKEAEKQGNVEVAVKAARVAVEAAKQAGDNDVLRKVAE QALEIAKKAAEQGDVGVMQKAMDVALRAAGQAG |
| | mScarlet-I_C | MGSKGEAVIKEFMRFKVHMEGSMNGHEFEIEGEGEGRPYEGTQT AKLKVTKGGPLPFSWDILSPQFMYGSRAFIKHPADIPDYYKQSFPE GFKWERVMNFEDGGAVTVTQDTSLEDGTLIYKVKLRGTNFPPDGP VMQKKTMGWEASTERLYPEDGVLKGDIKMALRLKDGGRYLADFKT |

| | | |
|---|---|---|
| | | TYKAKKPVQMPGAYNVDRKLDITSHNEDYTVVEQYERSEGRHSTGGMDELYKGSGSGDHALDRAALWALQAAIQAARLAAQAVLGTLQQPELFLQDLRSLVEAARILARLARQRGDLGGEEAREELREWLRELREALREAERVVAEHLSPVNLELALKALQILVNAAYVLAEIARDRGDEELLEAAAELAERAAEMAIRIAALAMEEGNFELALEALEIINEAARVLARIAHHRGNQELLEKAASLTHASAALSRAIAAILEGDVEKAVRAAQEAVKAAKEAGDNDMLRAVAIAALRIAKEAEKQGNVEVAVKAARVAVEAAKQAGDNDVLRLVSERALSIAASSVKQGNYEVKEKAIRVAKEANKQAG |
| | mNeonGreen_D | MGSKGEEDNMASLPATHELHIFGSINGVDFDMVGQGTGNPNDGYEELNLKSTKGDLQFSPWILVPHIGYGFHQYLPYPDGMSPFQAAMVDGSGYQVHRTMQFEDGASLTVNYRYTYEGSHIKGEAQVKGTGFPADGPVMTNSLTAADWCRSKKTYPNDKTIISTFKWSYTTGNGKRYRSTARTTYTFAKPMAANYLKNQPMYVFRKTELKHSKTELNFKEWQKAFTDVMGMDELYKGSGSGDHALDRAALWALQAAIQAARLAAQAVLGTLQQPELFLQDLRSLVEAARILARLARQRGDLGGEEAREELREWLRELREALREAERVVAEHLSPVNLELALKALQILVNAAYVLAEIARDRGDEELLEAAAELAERAAEMAIRIAALAMEEGNFELALEALEIINEAARVLARIAHHRGNQELLEKAASLTHASAALSRAIAAILEGDVEKAVRAAQEAVKAAKEAGDNDMLRAVAIAALRIAKEAEKQGNVEVAVKAARVAVEAAKQAGDNDVLKRVSETLLSIAAEATKQGNSEVMEKAIRVSEEAEKQAG |
| pD5 (in Figure 5; derived from pD5-14_rd47) | C (without mScarlet) | MGSDHALDRAALWALQAAIQAARLAAQAVLGTLQQPELFLQDLRSLVEAARILARLARQRGDLGGEEAREELREWLRELREALREAERVVAEHLSPVNLELALKALQILVNAAYVLAEIARDRGDEELLEAAAELAERAAEMAIRIAALAMEEGNFELALEALEIINEAARVLARIAHHRGNQELLEKAASLTHASAALSRAIAAILEGDVEKAVRAAQEAVKAAKEAGDNDMLRAVAIAALRIAKEAEKQGNVEVAVKAARVAVEAAKQAGDNDVLRLVSERALSIAASSVKQGNYEVKEKAIRVAKEANKQAG |
| | D (without mNeonGreen) | MGSDHALDRAALWALQAAIQAARLAAQAVLGTLQQPELFLQDLRSLVEAARILARLARQRGDLGGEEAREELREWLRELREALREAERVVAEHLSPVNLELALKALQILVNAAYVLAEIARDRGDEELLEAAAELAERAAEMAIRIAALAMEEGNFELALEALEIINEAARVLARIAHHRGNQELLEKAASLTHASAALSRAIAAILEGDVEKAVRAAQEAVKAAKEAGDNDMLRAVAIAALRIAKEAEKQGNVEVAVKAARVAVEAAKQAGDNDVLKRVSETLLSIAAEATKQGNSEVMEKAIRVSEEAEKQAG |
| | Neo-2/15_B | MPKKKIQLHAEHALYDALMILNIVKTNSPPAEEKLEDYAFNFELILEEIARLFESGDQKDEAEKAKRMKEWMKRIKTTASEDEQEEMANAIITILQSWIFSGGSGSGGGSGSAEWLERAAELALEVAERAARLAAEAWRTGVEEPRLVLRALENMVRAAHTLAEIARDNGERPDLEEIRERLEELAERLREELERAKRVAKEEEGKNLELALKALQILVNAAYVLAEIARDRGIELLLEAAAELAETAAELAIIAAAKAMEQGNLELALKALQIIVNAAYVLAEIARDRGNEELLEKAASLAEAAAALAEAIAAILEGDVEKAVRAAQEAVKAAKEAGDNDMLRAVAIAALRIAKEAEKQGNVEVAVKAARVAVEAAKQAGDNDVLRKVAEQALRIAKEAEKQGNVEVAVKAARVAVEAAKQAGDNDVLRKVAEQALEIAKKAAEQGDVGVMQKAMDVALRAAGQAG |
| | 4-1BB_mb1_B | MSGKATLEDLIALYEKGAAILEQIKPLVEKDMGLSNRTVATAIEEIKEAIKRVKKSGRIVYPIGLSIADNILALAQYYGNEKVAALAKELQKVGDAAAAVAEMVAAEEAGGSGSGGGSGSAEWLERAAELALEVAERAARLAAEAWRTGVEEPRLVLRALENMVRAAHTLAEIARDNGERPDLEEIRERLEELAERLREELERAKRVAKEEEGKNLELALKALQILVNAAYVLAEIARDRGIELLLEAAAELAETAAELAIIAAAKAMEQGNLELALKALQIIVNAAYVLAEIARDRGNEELLEKAASLAEAAAALAEAIAAILEGDVEKA |

| | | |
|---|---|---|
| | | VRAAQEAVKAAKEAGDNDMLRAVAIAALRIAKEAEKQGNVEVAVKAARVAVEAAKQAGDNDVLRKVAEQALRIAKEAEKQGNVEVAVKAARVAVEAAKQAGDNDVLRKVAEQALEIAKKAAEQGDVGVMQKAMDVALRAAGQAG |

**Supplementary Table 2. Cryo-EM data collection statistics for pD5-14.**

| | |
|---|---|
| **Data Collection** | |
| Microscope | Titan Krios (FEI) |
| Voltage (kV) | 300 |
| Detector | K3 (Gatan) |
| Energy Filter | BioQuantum Gif (Gatan) |
| Recording mode | Counting |
| Magnification | 105,000× |
| Movie micrograph pixel size (Å) | 0.843 |
| Dose rate ($e^-$/Å$^2$/s) | 11.31 |
| No. of frames per movie micrograph | 79 |
| Frame exposure time (ms) | 0.0505 |
| Movie micrograph exposure time (s) | 3.997 |
| Total dose ($e^-$/Å$^2$) | 45.21 |
| Under focus range (μm) | 0.8–1.8 |
| Total number of movies collected | 4871 |
| Total number of movies used | 4854 |
| **Map Processing** | |
| Extraction Box Size (pix) | 800 |
| Fourier crop to Box Size (pix) | 400 |
| Initial particle images (no.) | 640,897 |
| Final particle images (no.) | 209,004 |
| Map resolution (Å) | 4.30 |
| FCS threshold | 0.143 |
| Map resolution range (Å) | 3.71–5.28 |
| **Refinement** | |
| Initial model used | Design Model |
| Map resolution (Å) | 4.30 |
| FCS threshold | 0.143 |
| Model resolution range (Å) | 3.71–5.28 |
| Map sharpening B factor | 239.90 |
| Model composition | |
| Non-hydrogen atoms | 44,930 |
| Protein Residues | 9,060 |
| Ligands | N/A |
| B factors (Å) | |
| Protein | DeepEMhancer |
| Ligands | N/A |
| R.M.S. deviations | |
| Bond lengths (Å) (# > 4 σ) | 0.007 (0) |
| Bond angles (°) (# > 4 σ) | 1.471 (1) |
| Validation | |
| MolProbity score | 0.50 |
| Clashscore | 0.00 |
| Rotamer Outliers (%) | 0.00 |
| Ramachandran plot | |
| Favored (%) | 99.39 |
| Allowed (%) | 0.61 |
| Outliers (%) | 0.00 |

#### References


1. Goodsell, D. S. & Olson, A. J. Structural symmetry and protein function. *Annu. Rev. Biophys. Biomol. Struct.* **29**, 105–153 (2000).
2. Alberts, B. The Cell as a Collection of Protein Machines: Preparing the Next Generation of Molecular Biologists. *Cell* **92**, 291–294 (1998).
3. Lai, Y.-T., King, N. P. & Yeates, T. O. Principles for designing ordered protein assemblies. *Trends Cell Biol.* **22**, 653–661 (2012).
4. Khmelinskaia, A., Wargacki, A. & King, N. P. Structure-based design of novel polyhedral protein nanomaterials. *Curr. Opin. Microbiol.* **61**, 51–57 (2021).
5. Ueda, G. *et al.* Tailored design of protein nanoparticle scaffolds for multivalent presentation of viral glycoprotein antigens. *Elife* **9**, (2020).
6. Brouwer, P. J. M. *et al.* Enhancing and shaping the immunogenicity of native-like HIV-1 envelope trimers with a two-component protein nanoparticle. *Nat. Commun.* **10**, 4272 (2019).
7. Bruun, T. U. J., Andersson, A.-M. C., Draper, S. J. & Howarth, M. Engineering a Rugged Nanoscaffold To Enhance Plug-and-Display Vaccination. *ACS Nano* **12**, 8855–8866 (2018).
8. Rahikainen, R. *et al.* Overcoming Symmetry Mismatch in Vaccine Nanoassembly through Spontaneous Amidation. *Angew. Chem. Int. Ed Engl.* **60**, 321–330 (2021).
9. Boyoglu-Barnum, S. *et al.* Quadrivalent influenza nanoparticle vaccines induce broad protection. *Nature* **592**, 623–628 (2021).
10. Cohen, A. A. *et al.* Mosaic nanoparticles elicit cross-reactive immune responses to zoonotic coronaviruses in mice. *Science* **371**, 735–741 (2021).
11. Walls, A. C. *et al.* Elicitation of Potent Neutralizing Antibody Responses by Designed Protein Nanoparticle Vaccines for SARS-CoV-2. *Cell* **183**, 1367–1382.e17 (2020).
12. Walls, A. C. *et al.* Elicitation of broadly protective sarbecovirus immunity by receptor-binding domain nanoparticle vaccines. *Cell* **184**, 5432–5447.e16 (2021).
13. Marcandalli, J. *et al.* Induction of Potent Neutralizing Antibody Responses by a Designed Protein Nanoparticle Vaccine for Respiratory Syncytial Virus. *Cell* **176**, 1420–1431.e17 (2019).
14. Song, J. Y. *et al.* Safety and immunogenicity of a SARS-CoV-2 recombinant protein nanoparticle vaccine (GBP510) adjuvanted with AS03: A randomised, placebo-controlled, observer-blinded phase 1/2 trial. *eClinicalMedicine* **51**, (2022).
15. Liu, Y., Huynh, D. T. & Yeates, T. O. A 3.8 Å resolution cryo-EM structure of a small protein bound to an imaging scaffold. *Nat. Commun.* **10**, 1–7 (2019).
16. Liu, Y., Gonen, S., Gonen, T. & Yeates, T. O. Near-atomic cryo-EM imaging of a small protein displayed on a designed scaffolding system. *Proc. Natl. Acad. Sci. U. S. A.* **115**, 3362–3367 (2018).
17. Castells-Graells, R. *et al.* Cryo-EM structure determination of small therapeutic protein targets at 3 Å-resolution using a rigid imaging scaffold. *Proc. Natl. Acad. Sci. U. S. A.* **120**, e2305494120 (2023).
18. McConnell, S. A. *et al.* Designed Protein Cages as Scaffolds for Building Multienzyme Materials. *ACS Synth. Biol.* **9**, 381–391 (2020).
19. Divine, R. *et al.* Designed proteins assemble antibodies into modular nanocages. *Science* **372**, (2021).
20. Lutz, I. D. *et al.* Top-down design of protein nanomaterials with reinforcement learning. *bioRxiv* 2022.09.25.509419 (2022) doi:10.1101/2022.09.25.509419.
21. Mohan, K. *et al.* Topological control of cytokine receptor signaling induces differential effects in hematopoiesis. *Science* **364**, (2019).
22. King, N. P. *et al.* Computational design of self-assembling protein nanomaterials with atomic level accuracy. *Science* **336**, 1171–1174 (2012).
23. King, N. P. *et al.* Accurate design of co-assembling multi-component protein nanomaterials. *Nature* **510**, 103–108 (2014).
24. Hsia, Y. *et al.* Corrigendum: Design of a hyperstable 60-subunit protein icosahedron. *Nature* **540**, 150 (2016).
25. Bale, J. B. *et al.* Accurate design of megadalton-scale two-component icosahedral protein complexes. *Science* **353**, 389–394 (2016).
26. Wukovitz, S. W. & Yeates, T. O. Why protein crystals favour some space-groups over others. *Nat. Struct. Biol.* **2**, 1062–1067 (1995).
27. Kibler, R. D. *et al.* Design of pseudosymmetric protein hetero-oligomers. *Nat. Commun.* **15**, 1–12 (2024).
28. Lee, S. *et al.* Four-component protein nanocages designed by programmed symmetry breaking. *Nature* (2024) doi:10.1038/s41586-024-07814-1.
29. Dowling, Q. M. *et al.* Hierarchical design of pseudosymmetric protein nanocages. *Nature* (2024) doi:10.1038/s41586-024-08360-6.
30. Zhang, X., Fu, Q., Duan, H., Song, J. & Yang, H. Janus nanoparticles: From fabrication to (bio)applications.

*ACS Nano* **15**, 6147–6191 (2021).
31. Huehls, A. M., Coupet, T. A. & Sentman, C. L. Bispecific T-cell engagers for cancer immunotherapy. *Immunol. Cell Biol.* **93**, 290–296 (2015).
32. Li, X. *et al.* Preparation and application of Janus nanoparticles: Recent development and prospects. *Coord. Chem. Rev.* **454**, 214318 (2022).
33. Zeng, Y. C. *et al.* Fine tuning of CpG spatial distribution with DNA origami for improved cancer vaccination. *Nat. Nanotechnol.* **19**, 1055–1065 (2024).
34. Jin, H., Cui, J. & Zhan, W. Enzymatic Janus liposome micromotors. *Langmuir* **39**, 4198–4206 (2023).
35. Lee, J., Sands, I., Zhang, W., Zhou, L. & Chen, Y. DNA-inspired nanomaterials for enhanced endosomal escape. *Proc. Natl. Acad. Sci. U. S. A.* **118**, e2104511118 (2021).
36. Hsia, Y. *et al.* Design of a hyperstable 60-subunit protein icosahedron. *Nature* **535**, 136–139 (2016).
37. Wang, J. Y. J. *et al.* Improving the secretion of designed protein assemblies through negative design of cryptic transmembrane domains. *Proc. Natl. Acad. Sci. U. S. A.* **120**, e2214556120 (2023).
38. Dauparas, J. *et al.* Robust deep learning-based protein sequence design using ProteinMPNN. *Science* eadd2187 (2022).
39. Lee, S. *et al.* Design of four component T=4 tetrahedral, octahedral, and icosahedral protein nanocages through programmed symmetry breaking. *bioRxiv* 2023.06.16.545341 (2023) doi:10.1101/2023.06.16.545341.
40. Jumper, J. *et al.* Highly accurate protein structure prediction with AlphaFold. *Nature* **596**, 583–589 (2021).
41. Grueninger, D. *et al.* Designed protein-protein association. *Science* **319**, 206–209 (2008).
42. Garcia-Seisdedos, H., Empereur-Mot, C., Elad, N. & Levy, E. D. Proteins evolve on the edge of supramolecular self-assembly. *Nature* **548**, 244–247 (2017).
43. Young, G. *et al.* Quantitative mass imaging of single biological macromolecules. *Science* **360**, 423–427 (2018).
44. Velas, L. *et al.* Three-dimensional single molecule localization microscopy reveals the topography of the immunological synapse at isotropic precision below 15 nm. *Nano Lett.* **21**, 9247–9255 (2021).
45. Kirichenko, E. Y., Skatchkov, S. N. & Ermakov, A. M. Structure and functions of gap junctions and their constituent connexins in the mammalian CNS. *Biochem. (Mosc.) Suppl. Ser. A Membr. Cell Biol.* **15**, 107–119 (2021).
46. Karatekin, E. *et al.* A 20-nm step toward the cell membrane preceding exocytosis may correspond to docking of tethered granules. *Biophys. J.* **94**, 2891–2905 (2008).
47. Watson, J. L. *et al.* De novo design of protein structure and function with RFdiffusion. *Nature* **620**, 1089–1100 (2023).
48. Silva, D.-A. *et al.* De novo design of potent and selective mimics of IL-2 and IL-15. *Nature* **565**, 186–191 (2019).
49. Cao, L. *et al.* De novo design of picomolar SARS-CoV-2 miniprotein inhibitors. *bioRxivorg* (2020) doi:10.1101/2020.08.03.234914.
50. Gloegl, M. *et al.* Target-conditioned diffusion generates potent TNFR superfamily antagonists and agonists. *bioRxiv* (2024) doi:10.1101/2024.09.13.612773.
51. Dowling, Q. M. *et al.* Hierarchical design of pseudosymmetric protein nanoparticles. *bioRxiv* (2023) doi:10.1101/2023.06.16.545393.
52. Sosa, S. *et al.* Asymmetric bifunctional protein nanoparticles through redesign of self-assembly. *Nanoscale Adv.* **1**, 1833–1846 (2019).
53. Yang, E. C. *et al.* Computational design of non-porous pH-responsive antibody nanoparticles. *Nat. Struct. Mol. Biol.* **31**, 1404–1412 (2024).
54. Sheffler, W. *et al.* Fast and versatile sequence-independent protein docking for nanomaterials design using RPXDock. *PLoS Comput. Biol.* **19**, e1010680 (2023).
55. Padilla, J. E., Colovos, C. & Yeates, T. O. Nanohedra: using symmetry to design self assembling protein cages, layers, crystals, and filaments. *Proc. Natl. Acad. Sci. U. S. A.* **98**, 2217–2221 (2001).
56. de Haas, R. J. *et al.* Rapid and automated design of two-component protein nanomaterials using ProteinMPNN. *Proc. Natl. Acad. Sci. U. S. A.* **121**, e2314646121 (2024).
57. Meador, K. *et al.* A suite of designed protein cages using machine learning and protein fragment-based protocols. *Structure* **32**, 751–765.e11 (2024).
58. Huddy, T. F. *et al.* Blueprinting extendable nanomaterials with standardized protein blocks. *Nature* **627**, 898–904 (2024).
59. Lai, Y.-T. *et al.* Structure of a designed protein cage that self-assembles into a highly porous cube. *Nat. Chem.* **6**, 1065–1071 (2014).
60. Hoffnagle, A. M. & Tezcan, F. A. Atomically accurate design of metalloproteins with predefined coordination geometries. *J. Am. Chem. Soc.* **145**, 14208–14214 (2023).

61. Wicky, B. I. M. *et al.* Hallucinating symmetric protein assemblies. *Science* eadd1964 (2022).
62. Ellis, D. *et al.* Antigen spacing on protein nanoparticles influences antibody responses to vaccination. *Cell Rep.* **42**, 113552 (2023).
63. Pettersen, E. F. *et al.* UCSF ChimeraX: Structure visualization for researchers, educators, and developers. *Protein Sci.* **30**, 70–82 (2021).
64. Meng, E. C. *et al.* UCSF ChimeraX: Tools for structure building and analysis. *Protein Sci.* **32**, e4792 (2023).
65. Goddard, T. D. *et al.* UCSF ChimeraX: Meeting modern challenges in visualization and analysis. *Protein Sci.* **27**, 14–25 (2018).
66. Sanchez-Garcia, R. *et al.* DeepEMhancer: a deep learning solution for cryo-EM volume post-processing. *bioRxiv* (2020) doi:10.1101/2020.06.12.148296.
67. Liebschner, D. *et al.* Macromolecular structure determination using X-rays, neutrons and electrons: recent developments in Phenix. *Acta Crystallogr. D Struct. Biol.* **75**, 861–877 (2019).
68. Davis, I. W. *et al.* MolProbity: all-atom contacts and structure validation for proteins and nucleic acids. *Nucleic Acids Res.* **35**, W375–83 (2007).
69. Kidmose, R. T. *et al.* Namdinator - automatic molecular dynamics flexible fitting of structural models into cryo-EM and crystallography experimental maps. *IUCrJ* **6**, 526–531 (2019).
70. Croll, T. I. ISOLDE: a physically realistic environment for model building into low-resolution electron-density maps. *Acta Crystallogr. D Struct. Biol.* **74**, 519–530 (2018).
71. Emsley, P., Lohkamp, B., Scott, W. G. & Cowtan, K. Features and development of coot. *Acta Crystallogr. D Biol. Crystallogr.* **66**, 486–501 (2010).
72. wwPDB consortium. Protein Data Bank: the single global archive for 3D macromolecular structure data. *Nucleic Acids Res.* **47**, D520–D528 (2019).
73. Ellis, D. *et al.* Structure-based design of stabilized recombinant influenza neuraminidase tetramers. *Nat. Commun.* **13**, 1825 (2022).
